# A dual allosteric pathway drives human mitochondrial Lon

**DOI:** 10.1101/2021.06.09.447696

**Authors:** Genís Valentín Gesé, Saba Shahzad, Carlos Pardo-Hernández, Anna Wramstedt, Maria Falkenberg, B. Martin Hällberg

## Abstract

The hexameric, barrel-forming, AAA+ protease Lon is critical for maintaining mitochondrial matrix protein homeostasis. Efficient substrate processing by Lon requires the coordinated action of six protomers. Despite Lon’s importance for human health, the molecular bases for Lon’s substrate recognition and processing remain unclear. Here, we use a combination of biochemistry and electron cryomicroscopy (cryo-EM) to unveil the structural and functional basis for full-length human mitochondrial Lon’s degradation of mitochondrial transcription factor A (TFAM). We show how opposing protomers in the Lon hexamer barrel interact through their N-terminal domains to give what resembles three feet above the barrel and help to form a triangular pore located just above the entry pore to the barrel. The interactions between opposing protomers constitute a primary allosteric regulation of Lon activity. A secondary allosteric regulation consists of an inter-subunit signaling element in the ATPase domains. By considering the ATP or ADP load in each protomer, we show how this dual allosteric mechanism in Lon achieves coordinated ATP hydrolysis and substrate processing. This mechanism enforces sequential anti-clockwise ATP hydrolysis resulting in a coordinated hand-over-hand translocation of the substrate towards the protease active sites.

## Introduction

Deficiencies in mitochondrial function are linked to multiple diseases and age-related pathologies such as cancer, metabolic syndromes, inflammatory responses, and neurodegenerative disorders (Baker et al., 2011). The mitochondrial protein quality control (PQC) system maintains optimal mitochondrial function (Baker et al. 2014), with several key proteases that play fundamental roles (Puchades et al., 2020, Sauer & Baker, 2011). These include the mitochondrial matrix protease Lon, which is an AAA+ (ATPase Associated with various cellular Activities) protease. Mutations in the Lon gene cause Cerebral, Ocular, Dental, Auricular Skeletal (CODAS) syndrome, a complex multisystemic and developmental disorder, as well as of other mitochondrial-associated disease presentations (Peter et al., 2018; Strauss et al., 2015). Lon prevents the accumulation of toxic aggregates by selectively degrading proteins that are destabilized due to mutations or oxidative stress resulting from oxidative phosphorylation (Pinti et al., 2016; Quirós et al., 2014). In addition, Lon regulates mitochondrial gene expression through at least two mechanisms. First, the mitochondrial transcription factor A (TFAM), which is critical for mitochondrial DNA maintenance and expression, is specifically degraded by Lon (Lu et al., 2013). Secondly, upon stress conditions that trigger the mitochondrial unfolded protein response (UPR), Lon specifically degrades MRPP3, the active nuclease subunit in mitochondrial RNase P that performs the initial step in mitochondrial RNA processing (Münch & Harper, 2016; Reinhard et al., 2017).

A human Lon monomer contains a mitochondrial targeting sequence, a Lon-associated N-terminal (LAN) domain, an α-helical neck region, an AAA+ domain, and a protease domain (PD) in a single chain. This contrasts with other AAA+ proteases such as ClpXP or the 26S proteasome (Baker & Sauer, 2012; Bard et al., 2018; Lee et al., 2011), in which the ATPase and PD domains are in separate chains and function as obligate heterooligomers. Six Lon monomers assemble into a barrel-shaped structure (Cha et al., 2010; Duman & Löwe, 2010; Kereïche et al., 2016; Vieux et al., 2013; Shin et al., 2020; Shin et al., 2021; Zhang et al., 2020). The LAN domains specifically recognize and capture fully or partially unfolded substrates (Tzeng et al., 2021; Wohlever et al., 2014), which are then threaded towards the barrel chamber for degradation using the chemical energy from ATP hydrolysis (Duman & Löwe, 2010; He et al., 2018; Li et al., 2005, 2010; Vieux et al., 2013). Moreover, the LAN domains together with the neck region play a critical role in oligomerization (Kereïche et al., 2016). The unfolded substrate is threaded through the Lon barrel by two flexible loops in the AAA+ domain, namely pore loop 1 and 2, towards the protease domain at the bottom (Shin et al., 2020; Shin et al., 2021; Zhang et al., 2020). The protease active site is formed by a Ser-Lys dyad that is guarded by a loop (termed gate loop) that opens upon Lon hexamerization (Botos et al., 2004; Lin et al., 2016).

In this work, we set out to investigate the structural elements in Lon that coordinate ATP hydrolysis, substrate threading, and degradation. To this end, we determined cryo-EM single-particle structures of several different variants and catalytic states of human mitochondrial Lon.

## Results

### Structure of substrate-engaged human mitochondrial Lon

We cryogenically trapped the full-length mature form of human mitochondrial Lon (68–959; (Vaca Jacome et al., 2015) during active TFAM degradation (30 s incubation) to determine the structure of substrate-engaged Lon (Lon^SE^) using single-particle cryo-EM (Fig. 1A–C; Fig. S1). Human Lon forms a hexamer with overall dimensions 200 × 120 × 120 Å. In our reconstruction the maps for the AAA+ and protease domains were of excellent quality, thus enabling molecular model building. We observed density for a cryogenically trapped substrate peptide in the neck region and the pore of the Lon-hexamer barrel (Fig. 1B). Focused refinement and data collection using a tilted microscope stage enabled us to reveal the arrangement of the LAN domains and the neck region of Lon (Fig. 1C). Here, the reconstructed density enabled modelling of secondary-structure elements in the neck region and rigid body docking of homology-modelled LAN domains. The six Lon-protomers (P1–P6) display in-plane six-fold symmetry for the protease domains (PDs) consistent with an active state of Lon while the AAA+ domains are tilted with respect to a central six-fold axis (Fig. 1C).

**Figure 1.**
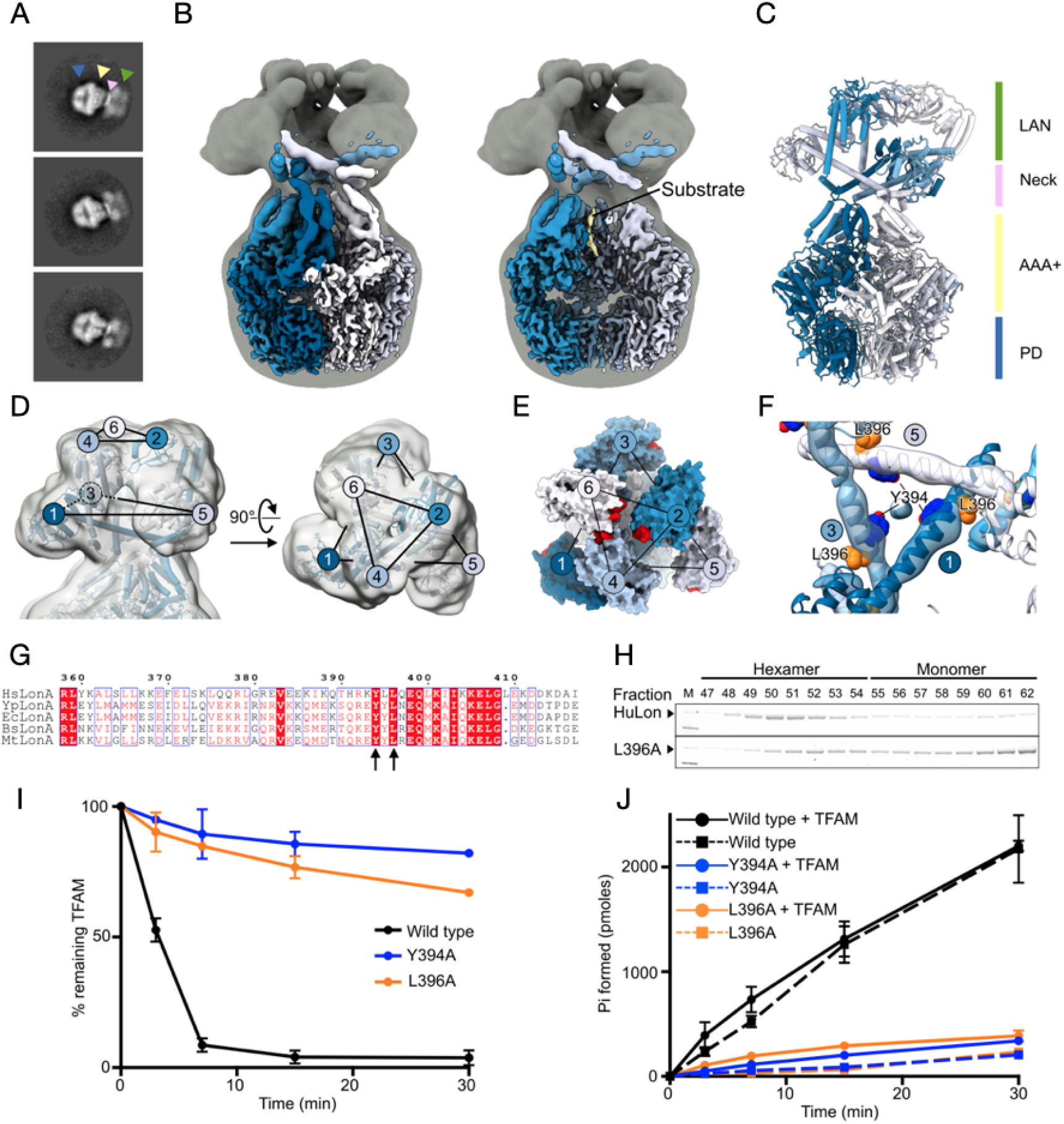
Overall structure of human Lon and the LAN domains/neck region. (**A**) Representative 2D class averages of Lon particles from cryo-EM of full-length Lon (68–959). The green, pink, yellow, and blue triangles indicate the LAN, the neck, the AAA+, and the PD regions, respectively. (**B**) Locally-filtered composite map of Lon^SE^ (left) and Lon^SE^ map after subtraction of protomers 5 and 6 to visualize the interior of the barrel (right), coloured from dark blue (protomer 1) to white (protomer 6, seam), with the substrate in yellow. The map is overlaid on a gaussian-filtered map to visualize the LAN domains (dark gray). (**C**) Overview of human Lon coloured from dark blue (protomer 1) to white (protomer 2). (**D**) Close-up view on the LAN and neck region of Lon^SE^. The electron density is shown as a grey transparent surface. Protomers 1–6 are coloured sequentially from blue to white. The stapled triangles formed by the LAN domains 1, 3, and 5 and by the domains 2, 4, and 6 are indicated with black lines. (**E**) View on the LAN domain region of Lon^SE^, shown as surface representation. The loops 127–129 and 146–151, which are shown in red, line the LAN domain pore and are important for substrate specificity as shown for *E. coli* Lon (Wohlever et al., 2014). (**F**) N-terminal pore formed by the neck helices in protomers 1, 3 and 5. Odd protomers Y394 (dark blue) and L396 (orange) are shown as spheres. (**G**) Multiple sequence alignment of the neck α-helical region (residues 358–417). The arrows indicate the conserved Y394 and L396. Bs, *Bacillus subtilis*; Ec, *Escherichia coli*; Hs, *Homo sapiens*; Mt, *Meiothermus taiwanensis*; Yp, *Yersinia pestis* (**H**) Gel filtration and SDS-PAGE analysis of the L396A Lon mutant shows impaired oligomerization. (**I**) Quantification of the proteolytic activity of Lon Y394A and L396A over time (0–30 min). Data is represented as mean ± standard deviation (n = 3). (**J**) Quantification of the ATP hydrolysis activity of Lon Y394A and L396A over time (0–30 min). Data is represented as mean ± standard deviation (n = 3).

The nucleotide state in each protomer was assigned based on the electron density and the configuration of the catalytic site residues (Fig. S2A). P1 and P2 are ATP-bound. The catalytic residues of P3 are in line with an ATP-bound state, but a weaker γ-phosphate density indicates ongoing hydrolysis in the cryogenically trapped active complex (Fig. S2B). We therefore assign the P3 Lon^SE^ protomer as ATP’. Lon^SE^ P4 and P5 are ADP-bound, while the “seam” protomer P6 is ATP-bound. For the Lon^SE^ protomers with bound ATP/ATP’ (P1–P3, P6) the AAA+-domains are located further from their respective PD in comparison to the AAA+-domains in the protomers with bound ADP (Fig S2C).

### The neck region supports oligomerization and interprotomer communication

The LAN domains form trimer-of-dimers tripod-like structures above the Lon barrel (Fig. 1D). We modelled the LAN dimers based on the highest excluded surface-area crystal contacts observed in earlier crystal structures of LAN domain homologs (Li et al., 2005; Li et al., 2010, Chen et al., 2019).

The best fitting dimer was used as a template for homology modelling (PDB 3LJC, Li et al., 2010) and the resulting model was rigid body fitted in triplicate. The LAN domains of opposing protomers in the Lon hexamer interact to form three feet (i.e., LAN P1 interacts with LAN P4, LAN P2 with LAN P5 and LAN P3 with LAN P6) (Fig. 1D). The LAN domains form a narrow pore that is lined by the loops 127–129 and 146–151 (Fig 1E). The α-helices in the neck of P1, P3, and P5 point towards the central axis and form a triangular pore (neck pore), which is ~6 Å wide, aligns with the central axis of the Lon hexamer, and is located ~20 Å above the ATPase domains (Fig 1F). The neck α-helices of P2, P4, and P6 are kinked by approximately 60°, point away from the central axis, and wrap around the opposing protomer α-helix, while the three LAN domains come together at the centre. At the same time, the α-helices form a knot that restrains the movement of the AAA+ domains during ATP hydrolysis (Fig. 1F). This knot can distribute the mechanical force generated by ATP binding and hydrolysis in each protomer to the neck and LAN domain regions of all protomers. In the neck of Lon, L396, in the interface between opposing neck helices, and Y394, which lines the neck pore, are both conserved (Fig. 1 F–G). A L396A mutation perturbs the interface between opposing helices and thereby impairs hexamerization (Fig 1H). Consequently, the L396A mutation causes a decrease in the protease and ATPase activity. Furthermore, a Y394A mutation in the neck pore severely impairs both TFAM degradation and ATP hydrolysis (Fig. 1 I–J and Table S1). These results highlight the importance of the neck region for interprotomer coordination, substrate recognition, and oligomerization.

### The pore loops contacting the substrate form a helix around the central axis

The Lon^SE^ map shows additional density in the barrel pore that we attributed to a substrate peptide (Fig 1B; Fig 2). The substrate side chains are not resolved, probably due to averaging different substrates during single-particle reconstruction, and therefore a polyalanine sequence was modelled. The substrate is contacted by two pore loops (termed pore loop 1 and 2) in P1–P3 but is contacted only by pore loop 1 in P4 and pore loop 2 in P5. The substrate is not contacted at all by P6 (seam protomer, Fig 2A). The final modelled substrate is 11 residues long and adopts a twisted β-strand conformation that follows the right-handed helical arrangement of the pore loops.

**Figure 2.**
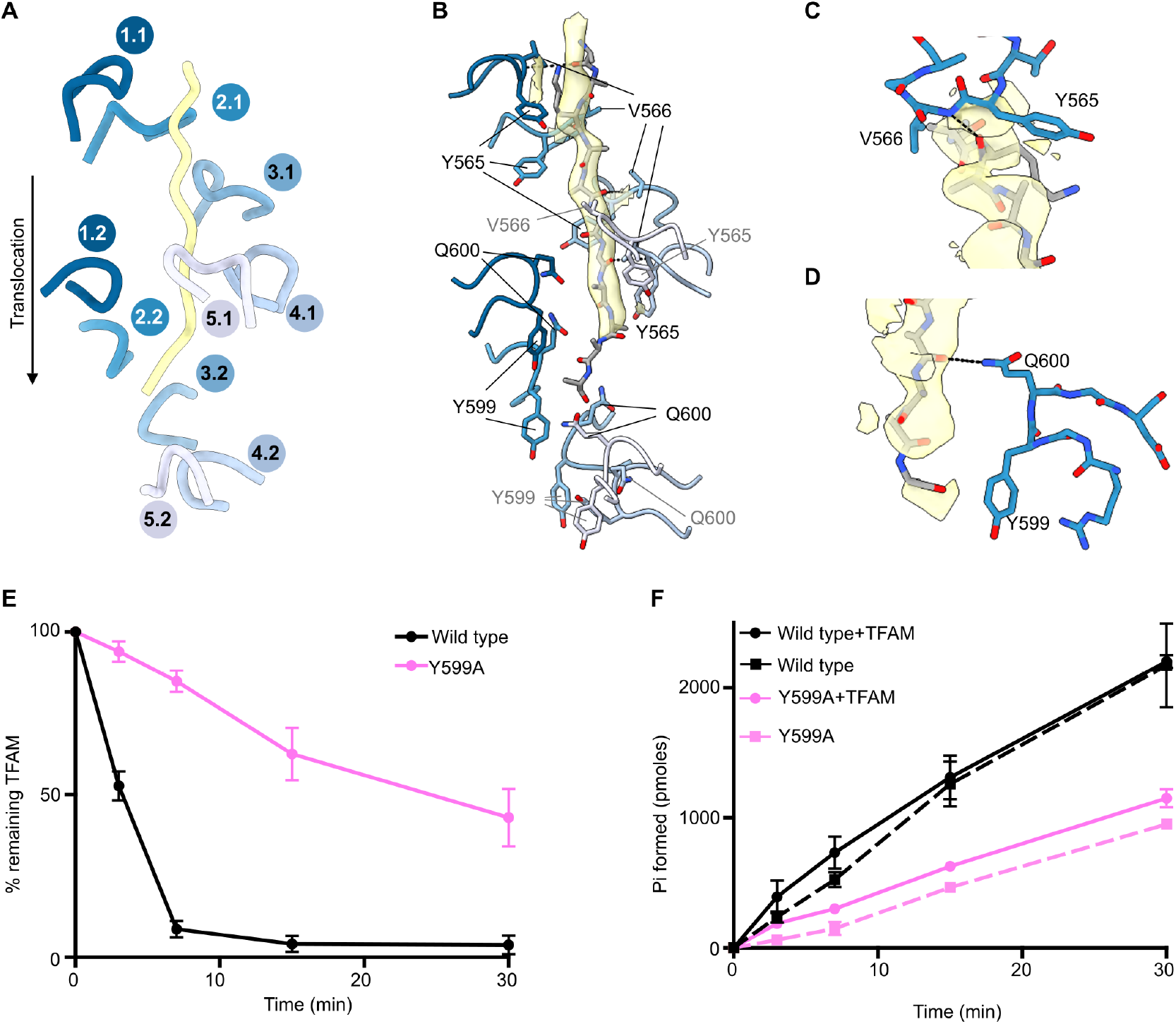
Lon interaction with the substrate. (**A**) The pore loops 1 (1.1–5.1) and 2 (1.2–5.2), coloured from dark to light blue for P1–P5, adopt a spiral organization around the substrate (yellow). (**B**) Pore loops 1 and 2 (coloured from dark to light blue for P1–P5) shown in cartoon representation. Y565 and V566 for pore loop 1 and Y599 and Q600 for pore loop 2 are shown as sticks. Pore loop residues that do not interact with the substrate are labelled in grey. The substrate is shown in stick representation, with the density shown as a yellow transparent surface. (**C**) A representative pore loop 1 interaction with the substrate (P2, blue sticks). (**D**) A representative pore loop 2 interaction with the substrate (P2, blue sticks). The substrate density is shown as a yellow transparent surface. H-bonds are shown by black dashed lines. (**E**) Quantification of the proteolytic activity of the Lon Y599A mutant over time (0-30 min). Data is represented as mean ± standard deviation (n=3). (**F**) Quantification of the ATP hydrolysis activity of the Lon Y599A mutant over time (0-30 min). Data is represented as mean ± standard deviation (n = 3).

The bound substrate interacts in the same way with the pore loop 1 (Y565–V566) in P1–P4. First, the substrate backbone oxygen forms a hydrogen bond (H-bond) with the backbone nitrogen of V566. Then the first and second side chains C-terminal to the H-bond form van der Waals contacts with Y565 and V566. These interactions repeat every second substrate residue for P1–P4 (Fig 2B,C). The substrate interacts in a similar manner with the pore loop 2 of P1–P3 (Fig. 2B,D). Here, Q600 forms an H-bond with the substrate backbone (Fig. 2D). Furthermore, Y599 forms a π-π stacking interaction with the peptide bond of the substrate (Fig. 2D) and a Y559A mutation, that removes this interaction, significantly lessens substrate turnover capacity and diminishes Lon ATPase activity (Fig. 2E–F). In addition, based on the Cβ orientation, we expect side chains with polar substrates to interact with the pore loop 2 backbone. The substrate does not reach protomer 4 pore loop 2, but instead contacts the P5 Q600 via its C-terminal oxygen (Fig. 2B).

Taken together, the pore loops in Lon^SE^ have a right-handed staircase arrangement around the central vertical axis of the chamber. This supports the hand-over-hand model previously proposed for other AAA+ ATPases, for example the 26S proteasome (de la Peña et al., 2018), Hsp104 (Gates et al., 2017), YME1 (Puchades et al., 2017), Vps4-Vta1 (Monroe et al., 2017), VAT (Ripstein et al., 2017), ClpXP (Ripstein et al., 2020), and *Y. pestis* Lon (Shin et al., 2020), in which the substrate is transferred from one protomer to the next in a sequential manner. The pore loops have extensive secondary structure (β-hairpins) but are relatively mobile as judged from the weak electron density. We propose that they are a mobile lever arm, which threads the substrate towards the protease domain.

### Neighbouring protomers communicate via the ISS element

The allosteric coordination between protomers in AAA+ ATPases is typically enabled by contacts with the nucleotide in trans and non-conserved inter-subunit signalling elements. For example, classical AAA+ ATPases contain the ISS motif (DGF), which is important for allosteric interprotomer communication (Augustin et al., 2009). Lon belongs to the HCLR clade (named after the representative members HslU, ClpAB-D2, Lon, and RuvB) of AAA+ ATPases and, the sequence of the ISS motif is not conserved (Fig. 3A), but the corresponding region 612–DPEQ–615 in our Lon^SE^ structure nevertheless establishes interactions with the neighbouring protomer and the nucleotide in trans. This region is situated between the Walker B D590 and the sensor 1 N640, and comprises the pore loop 2 with an insertion that contacts the pore loop 1, PS1βH. This region occupies a strategic position for inter-subunit signalling and we refer to it as the ISS element. To better understand interprotomer communication in Lon we analysed the interactions established by the ISS element in the different nucleotide states.

**Figure 3.**
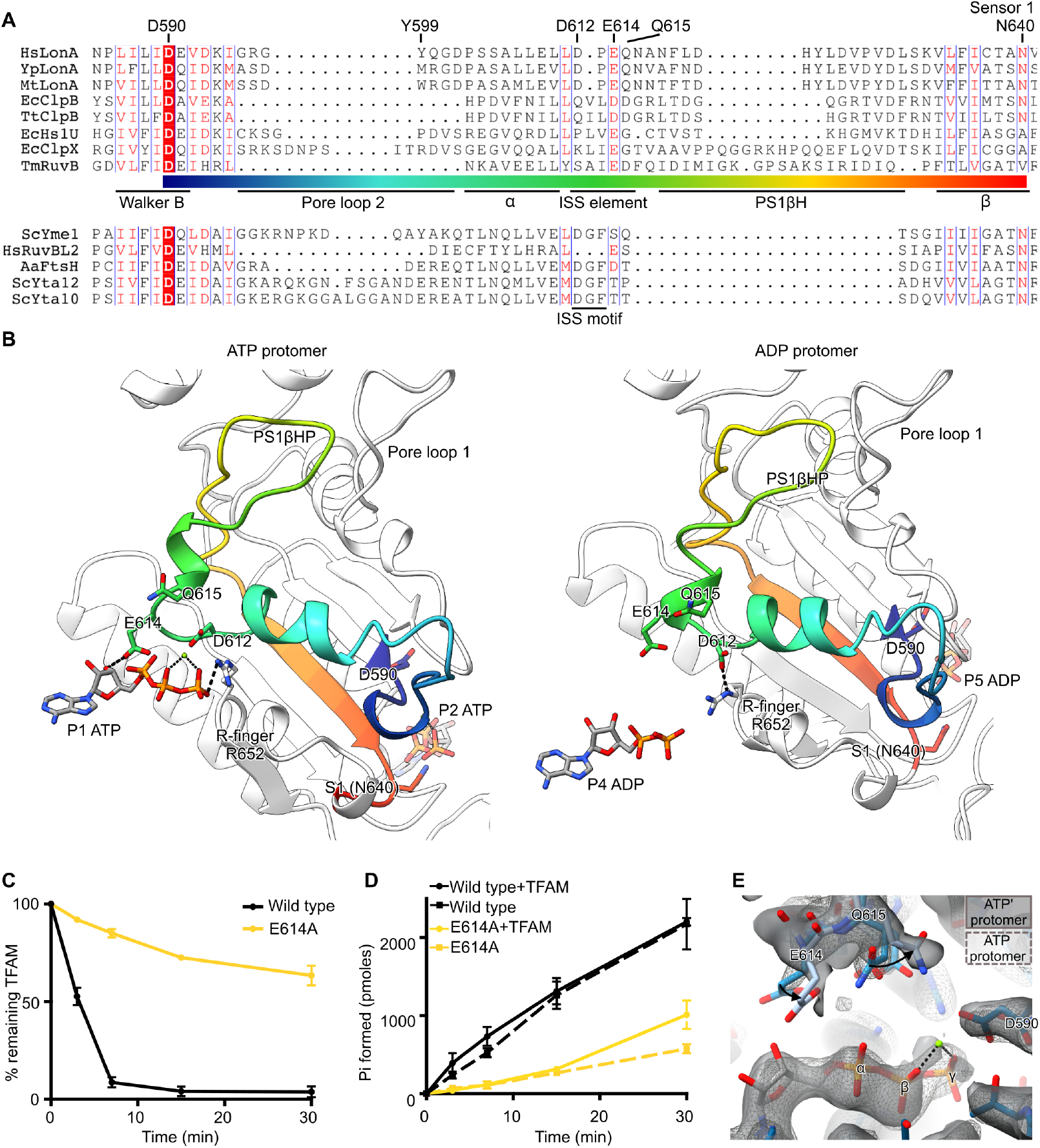
The ISS element. (**A**) Structure-based multiple sequence alignment of HCLR ATPases (upper block) and classic ATPases (lower block). Aa, *Aquifex aeolicus*; Ec, *Escherichia coli*; Hs, *Homo sapiens*; Mt, *Meiothermus taiwanensis*; Sc, *Saccharomyces cerevisiae*; Tm, *Thermotoga maritima*; Tt, *Thermus thermophilus*; Yp, *Yersinia pestis* (**B**) Cartoon representation of P2 (left) or P5 (right), with the region from D590 to N640 highlighted in the rainbow colours denoted in (A). The ISS element (D612, E614 and Q615), the R-finger (R652), the Walker B (D590), the sensor 1 (N640), as well as the nucleotide and the neighbouring nucleotide are shown as sticks. (**C**) Quantification of the proteolytic activity of the Lon AAA+ E614A mutant over time (0–30 min). Data is represented as mean ± standard deviation (n=3). (**D**) Quantification of the ATP hydrolysis activity of the Lon AAA+ E614A mutant over time (0–30 min). Data is represented as mean ± standard deviation (n = 3). (**E**) Comparison of the ISS element E614 and Q615 contacting the ATP and the ATP’ in trans, with the surrounding electron density shown as a mesh and as a semi-transparent surface, respectively. The surrounding P1 residues, ATP, and Mg^2+^ are also shown. The two protomers were superimposed on the ATP (P1) and ATP’ (P3) P-loops.

A key acidic residue in the ISS element is conserved in Lon (D612 in human Lon). This residue is also present in the ISS motif of classic AAA+ ATPases, where together with the R-finger (R652), they form a sensor-dyad that responds to the nucleotide state in the neighbouring protomer (Augustin et al., 2009). In accordance with this model, we here observe that in Lon P2–P4, D612 is ~3.2 Å from the R-finger, that interacts with the ATP-γ phosphate in trans. In P5–P1, the ISS element does not interact with the nucleotide and the R-finger moves even closer to D612 (Fig. 3B). Thus, D612 is important to convey a signal from the pore-loops or from ATP hydrolysis in cis to trigger ATP hydrolysis in trans.

A second acidic residue of the ISS element is conserved in the HCLR clade (E614 in human Lon, Fig. 3A). E614 is located at the hinge between the ISS motif and the PS1βH and interacts with the ribose of the neighbouring nucleotide (Fig. 3B,3E). Thus, E614 plays a structural role in positioning the ISS element and the PS1βH, and consequently a Lon E614A mutant has only 30% of the wild-type ATPase activity and seven-fold lower rate of TFAM hydrolysis (Fig. 3C,D and Table S1). In P3, which is undergoing hydrolysis, the neighbouring ISS element (from P4) moves closer to the nucleotide and the ISS Q615 interacts with the Walker B D590 (Fig. 3E). Given the important role of Walker B in catalysis, the ISS element has a direct role in catalysis via Q615. Notably, Q615 is not conserved in the HCLR clade and therefore is a Lon-specific allosteric residue (Fig. 3A). Taken together, the interactions established by the ISS element’s D612, E614, and Q615 influence the electrostatic charge distribution of the catalytic residues, keep catalysis under tight regulation, and behave as a precise relay that is sensitive to allosteric signals from the neighbouring protomer and the substrate.

### Substrate binding and hydrolysis in the Lon protease domain

To gain further insight into the regulatory mechanisms of Lon protease activity we determined the cryo-EM structure of a substrate-free form of Lon (Lon^Apo^, Fig. S3). We employed a fortuitously stable, catalytically inactive Lon mutant (containing the mutations G106P, R563W, and K594M), which also displays no proteolytic activity and limited ATPase activity (Fig S4 A-B). The Lon^Apo^ has a right-handed helical arrangement with a rise of approximately 8 Å (Fig. 4A, Fig. S4C) and all protomers are ADP-bound (Fig. S4D).

**Figure 4.**
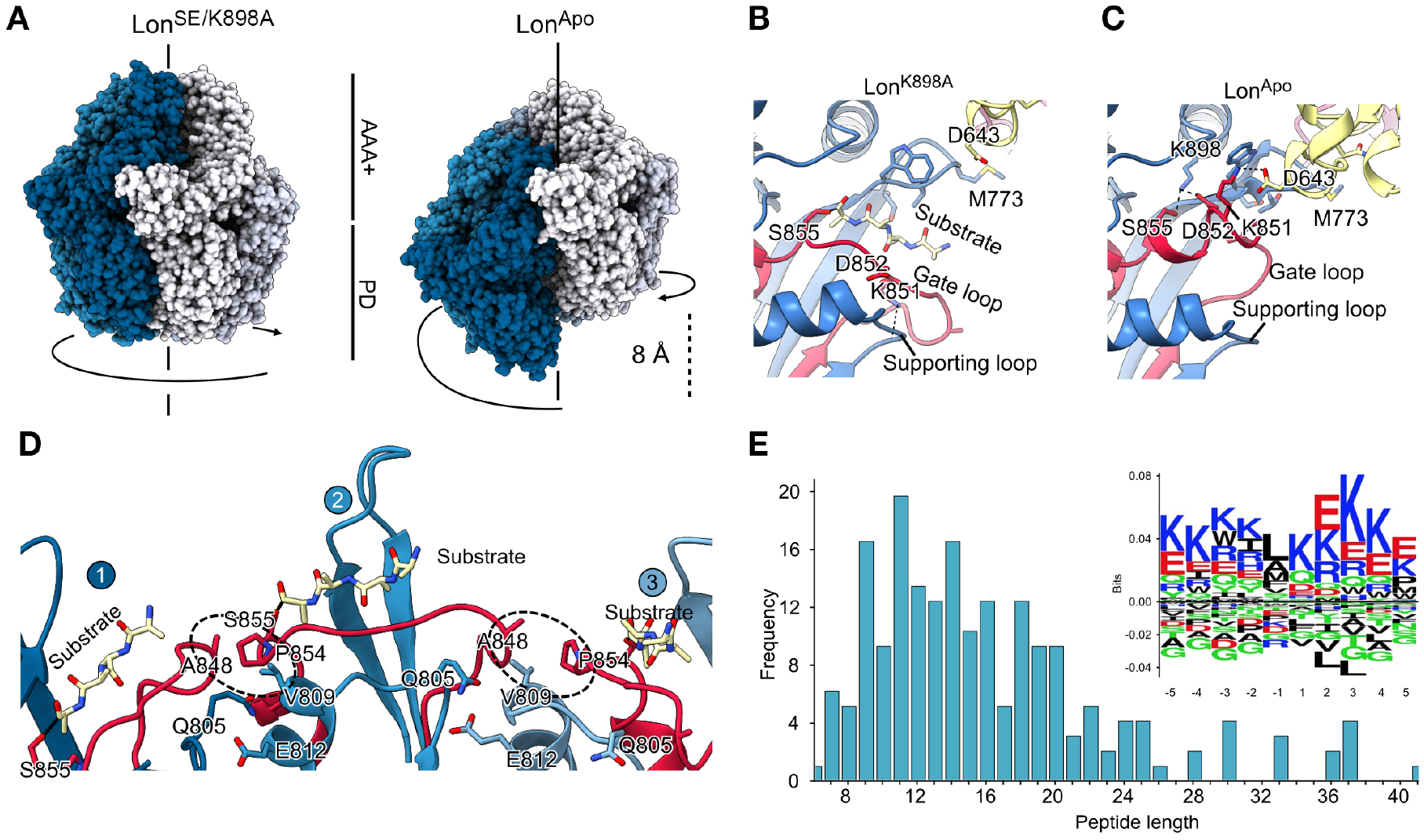
Structure and function of the protease domain. (**A**) Comparison of the protomer arrangement in Lon^Apo^ and Lon^SE/K898A^. Protomers 1–6 are coloured from blue to white. (**B**) Cartoon representation of Lon showing trapped substrate in Lon^K898A^ where the substrate is covalently bound to S855. The gate loop (red) is open in the protease active site in both Lon^SE^ and Lon^K898A^. The AAA+ domain is in yellow. (**C**) The gate loop in the closed conformation in Lon^Apo^. Relevant residues stabilizing the loop conformation and the catalytic dyad (S855, K898) are shown in stick representation. The substrate is shown as yellow sticks. (**D**) Substrate binding in the protease domain of Lon^K898A^ P1–P3, which are also representative for P4–P6. The substrate, in yellow sticks, adopts a β-strand conformation and augments the gate loop (red). The ester bond of the substrate to S855 is shown as a black line. The black dotted circle highlights the hydrophobic interaction. (**E**) Mass spectrometry analysis of the TFAM peptides produced by Lon. The histogram shows the peptide length frequency distribution. The inset shows the sequence logo generated from TFAM peptides aligned on the cleaved peptide bond and comprising residues before (−5 to −1) and after (1 to 5) the cleaved peptide bond.

Peptide bond hydrolysis is performed by the S855 and K898 catalytic dyad (Botos et al., 2004). Nucleophilic attack of S855 on the peptide bond creates an ester linkage between Lon and the substrate (Botos et al., 2004; Lee et al., 2007). The enzyme-substrate intermediate is resolved with the help of K898, which activates a water molecule for nucleophilic attack on the ester. Therefore, we introduced the mutation K898A (Lon^K898A^) in wt-Lon to trap the enzyme-substrate covalent intermediate of the hydrolysis reaction. We determined the structure of Lon^K898A^ in a similar manner to Lon^SE^ (Fig. S5) and the reconstruction revealed clear density for a trapped substrate in the protease domain. The trapped substrate is present in all six protomers, but its signal is approximately five-fold lower than that of the surrounding PD atoms. This suggests that only one of the six catalytic sites is occupied in each Lon particle. In addition, the substrate density level varies among the different protomers, indicating that the accessibility of the catalytic sites differs (Fig. S6A). We modelled tetra-alanine peptides in the maps and they show a β-strand conformation with the C-terminus of the substrate reaching S855 (Fig. 4B) and the N-terminus pointing towards the centre of the barrel.

The catalytic S855 is in an α-helix preceded by a loop (residues 844– 852, Fig. 4B) that functions as a gate controlling the accessibility to the catalytic site. In Lon^Apo^, this gate loop is in a closed conformation (Fig. 4C). The C-terminal part of the loop forms a 3_10_ helical turn, thereby extending the S855-containing α-helix. The turn is stabilized by a salt bridge between K851 and D643 in the α-subdomain. The 3_10_ helical turn positions the key residue D852 to form a salt-bridge with the catalytic K898 (Fig. 4C and Fig. S6B) and hinders the proteolytic activity. In contrast, the gate loop adopts an open conformation in Lon^SE^ and Lon^K898A^, in which the catalytic site is exposed (Fig. 4B). In this active conformation, the 3_10_ helix of the gate loop turns into a 3-residue β-strand that H-bonds to a supporting loop in the PD, comprising residues 803–805. The D852 in the β-strand is flipped 180°, thus breaking the interaction with the catalytic K898 and enabling proteolysis (Fig. 4B, and Fig. S6B). The β-strand also contains K851, which does not interact with the AAA+ domain as in the closed conformation but instead stabilizes the open conformation of the loop by interacting with the carbonyl oxygen of E846 in a preceding β-turn (Fig. S6B). The open loop conformation is further stabilized by hydrophobic interactions with the neighbouring protomer (Fig 4D). The substrate pairs in a parallel fashion with the gate loop and is guided to the catalytic centre (Fig 4D). In the main reconstructed conformation of Lon^K898A^ the P4 and P5 protease active sites are closest to the C-terminus of the substrate cryo-trapped in the pore loop (Fig. S6C).

The side chain preceding the cleaved peptide bond (−1 position) is accommodated in a hydrophobic pocket (Fig. S6D). This suggests that Lon preferentially cleaves substrates with a hydrophobic residue in the −1 position, and this is supported by MS analysis of peptides from Lon-mediated TFAM degradation (Fig. 4E, Fig. S7).

## Discussion

We determined high-resolution structures of substrate-engaged, substrate-trapped, and substrate-free forms of the human mitochondrial Lon protease. The substrate engaged (Lon^SE^) and substrate-trapped (Lon^K898A^) structures presented here were determined without using Walker-B mutations or artificial non-hydrolysable ATP-analogues typically utilized in similar studies of AAA-proteases. The even LAN domains (P2, P4, P6) form a narrow pore lined by the loops 127–129 and 146–151, which contribute to substrate recognition and capture. In the odd protomers (P1, P3, P5), the neck α-helices are straight, and the LAN domains are kept separate from each other (Fig. 1A–E). The neck α-helices in these protomers form a ~6 Å wide triangular pore. This neck pore aligns with the pore in the Lon barrel, indicating that Lon substrates are threaded through this neck pore. Accordingly, mutating the conserved Y394, which lines the neck pore, and L396, which stabilises the pore, impairs substrate degradation (Fig. 1F–J). The triangular organization of the even LAN domains is reminiscent of the ClpB N-terminal domains (Deville et al., 2017). In ClpB, three N-terminal domains form a hydrophobic pore aligned with the AAA+ barrel. The N-terminal domains are highly mobile and interact with the substrate to funnel it to the pore and similar to ClpB, the highly mobile even LAN domains function as clamps to capture folded, partially folded, or extended substrates and funnel them to the neck pore. The mechanical force of ATP hydrolysis is transferred to the LAN domains through the movement of the α-helices in the neck region while the movement of the neck α-helices threads the unfolded substrate through the neck pore towards the AAA+ domains, similarly to the thread guide in a sewing machine (Fig. 5).

**Figure 5.**
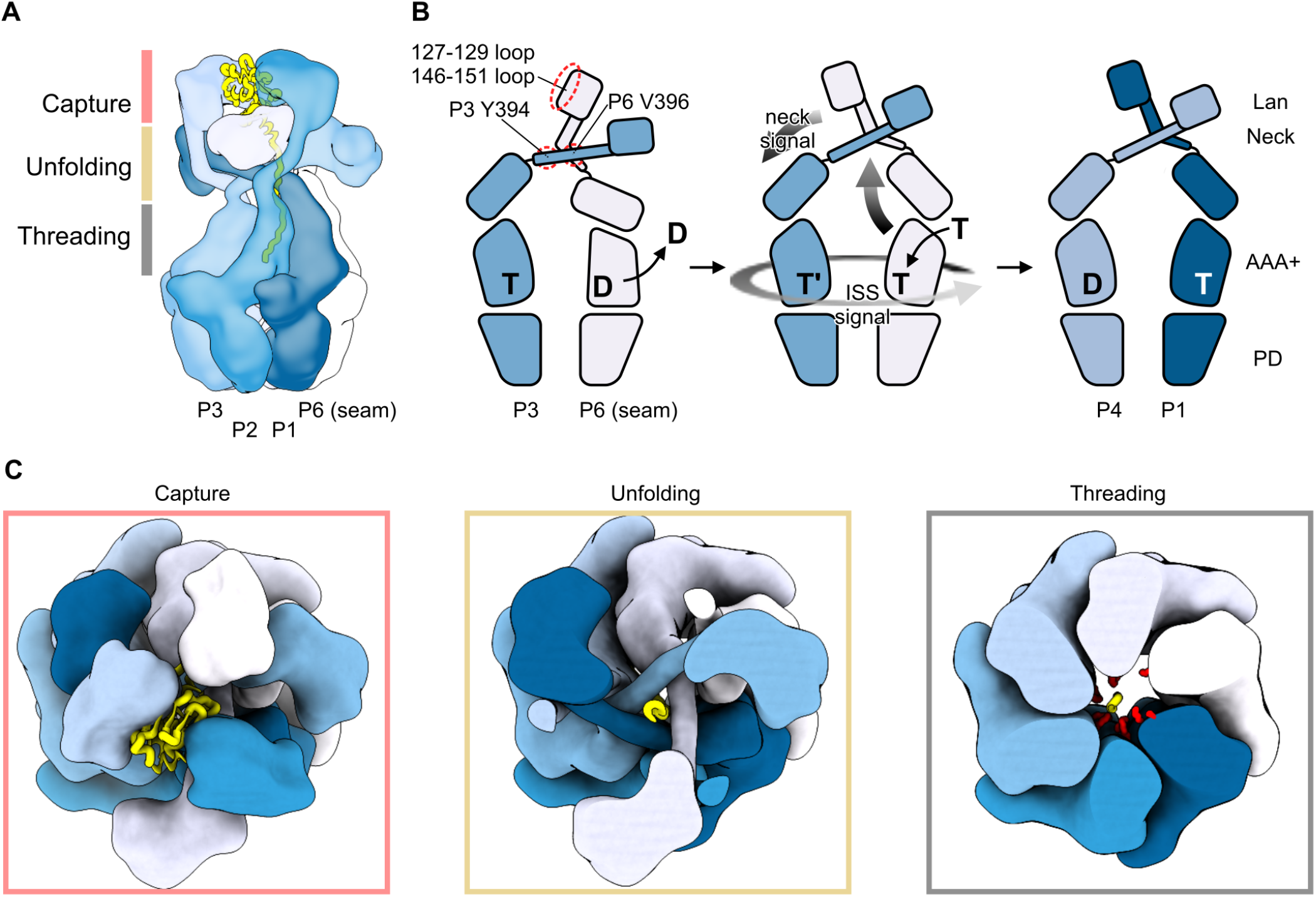
Allosteric mechanisms driving sequential ATP hydrolysis. (**A**) Lon in the process of degrading a substrate (yellow). The substrate is captured by the LAN domains, unfolded through the neck pore and threaded by the AAA+ domains towards the protease domains. (**B**) Cartoon representation of a pair of opposing protomers undergoing a cycle of ATP hydrolysis. An allosteric signal is transmitted across the barrel through the neck region upon nucleotide exchange at the seam protomer (P6), which then becomes P1. A second allosteric signal is transmitted around the barrel in the anti-clockwise direction via the ISS elements. This triggers ATP hydrolysis in P3, which then becomes P4. (**C**) Top view on the Lon hexamer (left), a cross-section at the neck pore level (middle) and at the AAA+ domain level (right). The pore loops are shown in red and the substrate in yellow.

In the AAA+ region, the substrate is translocated from one protomer to the next following a clockwise spiral staircase formed by the pore loops (what has been termed hand-over-hand translocation, Puchades et al., 2020). We can thus expect that the translocation rate in Lon is two residues per hydrolysed ATP. Pore loops 1 and 2 contain Y565 and Y599, respectively, that interact with the substrate backbone (Fig. 2A–C). Consistent with this observation, the mutation Y565H was identified in a patient with classical mitochondrial disease presentations, and impairs substrate binding and degradation (Peter et al., 2018). Although Y599 is not strongly conserved, physical and chemical functionality is retained with bulky hydrophobic side chains (M, F, or W) in Lon homologs and the mutation Y599A impairs substrate degradation as well as ATP hydrolysis (Fig. 2E-F).

Consistent with Lon^SE^ being in an active conformation, the PDs display in-plane six-fold symmetry and an open gate loop. Furthermore, reconstruction of covalently-trapped Lon^K898A^ obtained in the presence of substrate TFAM retains the in-plane six-fold symmetry of the PDs and shows substrate density in all protease domains. After the covalent bond between S855 and the TFAM C-terminus forms, substrate threading is stalled.

It is well established that ATP hydrolysis in the HCLR ATPases occurs in a coordinated manner. However, different models have been proposed depending on the different nucleotide loads observed in each protomer. In our comparison below it should be noted that our structure (Lon^SE^) is the only substrate-bound structure available for Lon without active-site mutations (Walker B) and without the use of non-hydrolyzable ATP analogs thereby making direct comparisons of nucleotide load difficult.

During preparation of this manuscript, a cryo-EM structure of a shortened human Lon construct (115–959) in an inactive conformation was presented (Shin et al., 2021). In that case, only the ATPase domains and PDs were ordered and the PDs were in a staggered non-active conformation. Part of the protein preparation contained a substrate presumably remnant from the overexpression host in the barrel pore (Shin et al., 2021). The structure contained ADP in the P6 and P5 protomers, and trapped ATP (or ATPγS) in the rest. Similarly, a *Y. pestis* Lon substrate-engaged structure was obtained with the help of a combination of ATPγS and Walker-B mutations (Shin et al., 2020). Based on these studies and other examples, it was proposed that ATP hydrolysis occurs in a sequential anti-clockwise order and that ADP to ATP exchange occurs in the seam position (Shin et al., 2020). However, in our Lon^SE^ structure the seam (P6) protomer is bound to ATP. The P5 and P4 protomers are loaded with ADP, while the P3 is primed for ATP hydrolysis considering the partial occupancy of the γ-phosphate and the configuration of the catalytic residues while P1 and P2 are loaded with ATP. Therefore, the Lon^SE^ structure presented here represents the stage immediately following nucleotide exchange in the seam protomer P6, and prior to its re-engagement in substrate binding. Therefore, ATP hydrolysis in P4 and priming in P3 should be concomitant with nucleotide exchange in the seam. This requires allosteric interprotomer communication between opposing protomers (seam (P6) and P3) through the neck region, which emerges as a new structural element supporting the sequential anti-clockwise mechanism of ATP hydrolysis and thereby constitutes a primary allosteric regulation (Fig 5 and Supplementary Movie S1).

A secondary allosteric regulation in human Lon is performed by the ISS element, that conveys signals from the pore-loops or from ATP hydrolysis in cis to trigger ATP hydrolysis in trans by influencing the electrostatic charge distribution of catalytic residues. Thereby the ISS elements helps to keep catalysis under tight regulation, and constitutes a precise relay sensitive to allosteric signals from the neighbouring protomer and the substrate. The role of PS1βH in this secondary allosteric regulation by the ISS element is highlighted by the S631F mutation, which is located at the base of PS1βH and is associated with the CODAS syndrome (Strauss et al., 2015). Hence, substrate degradation by human mitochondrial Lon follows a dual-allosteric pathway.

Understanding the Lon mechanism is important due to the multiple disease-causing mutations in Lon (Peter et al., 2018; Strauss et al., 2015), which are distributed over the entire sequence and impair its function in different ways (Fig S8). The structure of active human mitochondrial Lon presented here fosters further molecular understanding of human mitochondrial disease and may open the path to new therapeutic interventions.

## Materials and methods

### Production, expression and purification of Lon and TFAM

The gene for wild-type human mitochondrial LonP1 (UniProt - P36776) was obtained from the Mammalian Gene Collection (gi:21595090). The sequence for the mature protein (68-959) was PCR amplified and the purified PCR product was cloned into the pNIC28-Bsa4 vector (gi:124015065) that provides a 6x His-tag followed by a TEV protease cleavage site in the N-terminus. The resulting construct contained the spontaneously arisen mutations G106P, R563W and K594M. This construct was expressed, purified and employed in single-particle electron microscopy (as described below) resulting in the Lon^Apo^ structure. Site-directed mutagenesis (QuikChange-Lightning; Agilent) was used to revert the cloned sequence to the wild-type sequence. Mutations Y394A, L396A, Y599A or K898A were introduced in the wild-type sequence by site-directed mutagenesis.

For Lon and Lon mutants production, Rosetta™(DE3)pLysS competent cells (Novagen) were transformed and cultured in Terrific Broth media at 37°C for 5 h. Protein expression was induced with 1 mM IPTG at 16°C for 4 h. Cells were harvested by centrifugation, frozen in liquid nitrogen, thawed and scraped at 4°C in lysis buffer (25 mM Tris-HCl pH 8.0, 10 mM β-mercaptoethanol). Cells were then lysed using an Ultra-Turrax T3 homogenizer (IKA) and the lysate was incubated for 1 h with a previously equilibrated His-Select Nickel Affinity Gel (Sigma-Aldrich). Weakly-bound proteins were removed using buffer A (20 mM imidazole, 25 mM Tris-HCl pH 8.0, 400 mM NaCl, 10% glycerol, 10 mM β-mercaptoethanol), followed by elution with buffer B (250 mM imidazole, 25 mM Tris-HCl pH 8.0, 400 mM NaCl, 10% glycerol, 10 mM β-mercaptoethanol). Overnight dialysis in the presence of TEV protease was performed to remove the His tag and a new Nickel purification step was made as stated above, saving in this case the flow-through fraction. Proteins were subsequently purified over a 5 ml HiTrap Heparin column (GE Healthcare) and a 1 ml HiTrap Q column (GE Healthcare), both equilibrated in a buffer containing 20 mM Tris-HCl pH 8.0, 100 mM NaCl, 10% glycerol and 1 mM DTT. The columns were developed using a salt gradient (200 mM-1200 mM NaCl) in the equilibration buffer. Mature TFAM (amino acids 43–246) was cloned into the pJ411 vector (ATUM). The construct included a 6x His-tag followed by a TEV protease cleavage site in the N-terminus. Rosetta™(DE3)pLysS competent cells (Novagen) were transformed with the plasmid and cultured in Luria-Bertani Broth (LB) media at 37°C until reaching a OD_600_ of 0.6. Protein expression was induced with 1 mM IPTG at 16°C overnight. Protein purification followed the same steps as the mentioned above for Lon, with the absence of the HiTrap Q column. Protein purity was assessed by SDS-PAGE. Purified proteins were aliquoted and stored at −80°C.

### ATPase assay

In vitro ATPase assay was performed and quantified using the Malachite Green Phosphate Assay kit (BioAssay systems, POMG-25H). 0.5 μg of Lon (residues 68-959) wild-type and mutant versions were mixed, in presence or absence of 0.2 mg of TFAM, in a total volume of 15 μl per reaction with a buffer containing 50 mM Tris-HCl pH 8.0, 10 mM MgCl_2_, 0,1 mg/ml BSA, 2 mM ATP and 1 mM DTT. The reaction was started by addition of 2 mM ATP and incubated at 37°C for 0-30 min. Values at each sampled timepoint were calculated from three independent measurements. The inorganic phosphate released was calculated based on a absorbance standard curve established by 0-40 mM phosphate.

### TFAM proteolysis assay

In-vitro TFAM proteolysis assays were performed in 10 μl reaction volumes. 0.5 μg of Lon (residues 68-959) wild-type and mutant versions and 0.2 μg of TFAM were incubated at 37°C for 0-30 min in a buffer containing 50 mM Tris-HCl pH 8.0, 10 mM MgCl_2_, 0.1 mg/ml BSA, 2 mM ATP and 1 mM DTT. The reactions were stopped by the addition of SDS-PAGE loading buffer. Samples were run in a precast 4-20% gradient SDS-PAGE gel (BioRad, 567-8094). Relative band intensities were measured using ImageLab™ (BioRad) and calculations were made in order to provide % remaining TFAM values. Rate constants were obtained by fitting the data to y=a*(1-exp(-b*x)), where a is the maximum value of y and b is the apparent rate constant.

### Oligomerization assay/gel filtration analysis

Hexamerization of both wild-type and L396A Lon was tested by gel filtration chromatography using a Superose 6 Increase 3.2/300 column (GE Healthcare). The column was pre-equilibrated in running buffer (25 mM Tris–HCl, pH 8.0, 10% glycerol, 1 mM DTT, 50 mM NaCl and 2 mM ATP). The proteins were dialyzed into the corresponding running buffer and then incubated at 37°C for 10 min with 2 mM ATP and 10 mM MgCl_2_. The samples (200 μl) were injected onto the column through a loop. Fractions of 250 μl were collected and analysed by SDS-PAGE.

### Specimen preparation for cryo-EM

Ten microliters of 0.5–0.7 mg/ml of the different Lon constructs in buffer (20 mM HEPES pH 7.5, 100 mM NaCl, 10 mM MgCl_2_, 3% glycerol and 2 mM ATP) were added to 0.2 mg/ml TFAM and incubated for 30 s at room temperature. Immediately after, the samples (3.5 μl) were applied onto UltraAuFoil R 2/2 gold grids (Quantifoil Micro Tools GMBH; glow discharged with 25 mA for 60 s) at 4°C and 100% humidity in a Vitrobot Mk IV (Thermo Fisher Scientific). Sample excess were removed by blotting for 3 s using a blot force of 0, followed by vitrification in liquid ethane.

### Cryo-EM data acquisition and processing

The data were collected on a Krios G3i electron microscope (Thermo Fisher Scientific) at an operating voltage of 300 kV with Gatan BioQuantum K3 image filter/detector (operated with a 10 eV slit) at the Karolinska Institutet’s 3D-EM facility, Stockholm, Sweden. The data were collected using EPU (Thermo Fisher Scientific). An EFTEM SA magnification of 130,000x, was used resulting in a pixel size of 0.654 Å with a fluency of 51 e^−^/Å^2^ divided across 60 frames over 1.5 s. Target defocus values were set between −0.2 to −2 μm and using a stage tilt of 0°, −10° or −15°.

The data processing strategy is schematized in Figs S1, S3 and S5 for Lon^SE^, Lon^Apo^ and Lon^K898A^, respectively. For all data collections, motion correction, initial CTF estimation and particle picking was performed in Warp (Tegunov & Cramer, 2019), using a pre-trained deep-learning-based particle picking algorithm (BoxNet2Mask_20180918 model).

For Lon^SE^, the particles were extracted using a 600-pixel box (0.654 Å/ pix) and imported into CryoSPARC v2.15 (Punjani et al., 2017). Ab-initio reconstruction was performed with all particles using 4 classes, followed by heterogeneous refinement of the 4 classes. One of the classes yielded a high-resolution Lon reconstruction (179,823 particles). The particles from this class were filtered by 2D classification (156,036 particles) and subjected to homogeneous refinement. The particles were then imported into RELION 3.1 for Bayesian polishing (Scheres, 2012; Zivanov et al., 2019). The polished particles were re-imported into CryoSPARC v2.15 for local refinement with either a protease domain or ATPase focus. To improve the map of the N-terminal region, the particles were down sampled to a 200-pixel box (1.962 Å/pix) and classified without particle alignments using a mask on the N-terminal region in RELION 3.1. Two of the classes showed a higher detail in the N-terminal region (55,952 particles). These classes were pooled, re-extracted into a 400-pixel box (1.308 Å/pix) and re-imported into CryoSPARC v2.15 for homogeneous refinement followed by local refinement with a focus on the N-terminal region. As a final step, the three focused maps were combined using the ‘Combine focused maps’ tool in PHENIX (Liebschner et al., 2019).

For Lon^Apo^, the data were further processed in CryoSPARC v2.15. Particles were extracted to a 400-pixel box (1.308 Å/pix). After 2D classification, the manually selected good 2D classes (143,637 particles) were used for ab-initio reconstruction using 4 classes. The 4 classes were used as 3D volume templates in heterogeneous refinement of the entire particle set (349,068 particles). One of the classes resulted in a good Lon reconstruction (118,890 particles) and the particles were thereafter used in homogeneous refinement followed by a local refinement with a mask covering the ATPase and protease domains.

For Lon^K898A^, the particles were extracted using a 600-pixel box (0.654 Å/ pix). The particles were filtered by 2D classification in CryoSPARC v2.15, and the manually selected good particles were used for ab-initio reconstruction in 4 classes. The 4 classes were input into heterogeneous refinement. Two of the classes produced high resolution reconstructions of Lon. The particles from these two classes were pooled and subjected to homogeneous refinement. The particles were then imported into RELION 3.1 for Bayesian polishing. The polished particles were input into homogeneous refinement followed by local refinement with a mask on the ATPase or protease domains in CryoSPARC v2.15. The map was post-processed by LocScale (Jakobi et al., 2017) using the Lon^K898A^ atomic coordinates, which had been previously refined against the local refinement map. All reported resolutions were estimated using gold-standard Fourier shell correlation coefficient at a level of 0.143 (Rosenthal & Henderson, 2003). Please see table S2 for further data collection and processing information.

### Model building and refinement

The initial model of Lon^Apo^ was derived by homology modelling in SWISS-MODEL (Arnold et al., 2006) using the crystal structure of *B. subtilis* Lon (3M6A with 415-948 amino acids) as a template, with which it has 47% identity. Each protomer was rigid body fitted into the cryo-EM density map using ChimeraX (Goddard et al., 2018) and was used as a starting model for manual building in Coot v0.9 (Emsley et al., 2010). Multiple rounds of real-space refinement in PHENIX (Liebschner et al., 2019), reciprocal space refinement in REFMAC5 (Murshudov et al., 2011) and manual rebuilding with Coot were performed. The model was validated using Molprobity (Chen et al., 2010) and EMRinger (Barad et al., 2015).

The Lon^SE^ model was built from Lon^Apo^. First, the six protomers were individually rigid body fitted into the density using Chimera. The docked template was then subjected to real space refinement in Coot v0.9 using PROSMART-generated (Nicholls et al., 2014) self-restraints. To model the substrate in the AAA+ domain, a β-strand of 11 residues was built de novo in the density. As the side chains are not visible, all residues were modelled as Ala. The model was further refined by using multiple cycles of PHENIX real space refine, REFMAC5 and Coot.

For Lon^K898A^, the substrates in the protease domains were built de novo as tetra-alanine β-strands covalently linked in the C-terminus to S855. The model was refined using PHENIX real space refine.

The density for the N-terminal region reveals that the LAN domains form a trimer of dimers. We analysed the available crystal structures of E. coli LAN domains and extracted all possible dimers, considering both dimers within the asymmetric unit as well as crystallographic dimers. The best fitting structure came from a crystallographic dimer in the 3LJC structure (Li et al., 2010). The dimer was used as a template for homology modelling of the human Lon N-terminal region using SWISS-MODEL. The model was idealised using REFMAC5 and fitted in triplicate using ChimeraX. The neck helix in protomers 2, 4 and 6 was bent by approximately 60° in order to connect to the corresponding ATPase domain. The LAN domains were rigid body refined against the N-terminal focused map. Local resolution maps were computed using CryoSparc v2.15. Measurements and figures of the fitted model and density maps were made using ChimeraX. Please see table S3 for further model building, refinement and validation information.

## Acknowledgements

This work was supported by the Swedish Research Council (2018-02439 to MF and 2018-3808 to BMH), the Knut and Alice Wallenberg Foundation (KAW 2017.0080 to MF and BMH) and Wallenberg Scholar (to MF), the Swedish Cancer Foundation (2019-816 to MF) and the European Research Council (2016-683191 to MF). CP was supported by the Marie Sklodowska-Curie ITN-REMIX (721757). We acknowledge M. Carroni, M. Hall and J. Conrad for support in preliminary cryo-EM screening and collection at the SciLifeLab cryo-EM facilities funded by the Knut and Alice Wallenberg, Family Erling Persson, and Kempe foundations. All final cryo-EM data were collected at the Karolinska Institutet’s 3D-EM facility.

## Author contributions

GVG, SS and BMH performed cryo-EM sample preparation, data collection, processing, model building and refinement. CP and AW produced the protein expression constructs. CP performed protein expression, purification and biochemical assays. GVG, CP, SS and BMH wrote the manuscript, with input from all authors. MF and BMH supervised the research.

## Data availability

Structures and maps presented in this paper have been deposited to the Protein Data Bank (PDB) and Electron Microscopy Data Base (EMDB) with the following accession codes: Lon^SE^ main composite map and three focused refinement maps EMD-13146, the Lon^SE^ atomic coordinates PDB ID: 7P09, Lon^SE^ atomic coordinates comprising the LAN domains and neck region, which due to the lower resolution were homology modelled and docked as rigid bodies GitHub (https://github.com/genisvalentin/hsLonP1.git); Lon^Apo^ map EMD-13147; Lon^Apo^ atomic coordinates PDB ID: 7P0B; Lon^K898A^ map EMD-13148; Lon^K898A^ atomic coordinates PDB 7P0M.

## References

Arnold, K., Bordoli, L., Kopp, J., & Schwede, T. (2006). The SWISS-MODEL workspace: A web-based environment for protein structure homology modelling. Bioinformatics, 22(2), 195–201

Augustin, S., Gerdes, F., Lee, S., Tsai, F. T. F., Langer, T., & Tatsuta, T. (2009). An Intersubunit Signaling Network Coordinates ATP Hydrolysis by m-AAA Proteases. Mol Cell, 35(5), 574–585

Baker, M. J., Palmer, C. S., & Stojanovski, D. (2014). Mitochondrial protein quality control in health and disease. Br J Pharmacol. 171(8):1870–1889

Baker, M. J., Tatsuta, T., & Langer, T. (2011). Quality control of mitochondrial proteostasis. Cold Spring Harb Perspect Biol. 3(7):a007559

Baker, T. A., & Sauer, R. T. (2012). ClpXP, an ATP-powered unfolding and protein-degradation machine. Baker TA, Sauer RT. Biochim Biophys Acta. 1823(1):15–28.

Barad, B. A., Echols, N., Wang, R. Y. R., Cheng, Y., Dimaio, F., Adams, P. D., & Fraser, J. S. (2015). EMRinger: Side chain-directed model and map validation for 3D cryo-electron microscopy. Nat Methods, 12(10), 943–946

Bard J. A. M., Goodall, E. A., Greene, E. R., Jonsson, E., Dong, K. C., & Martin, A. (2018). Structure and Function of the 26S Proteasome. Annu Rev Biochem. 87:697–724.

Botos, I., Melnikov, E. E., Cherry, S., Tropea, J. E., Khalatova, A. G., Rasulova, F., Dauter, Z., Maurizi, M. R., Rotanova, T. v, Wlodawer, A., & Gustchina, A. (2004). The catalytic domain of Escherichia coli Lon protease has a unique fold and a Ser-Lys dyad in the active site. J Biol Chem, 279(9), 8140–8148

Cha, S. S., An, Y. J., Lee, C. R., Lee, H. S., Kim, Y. G., Kim, S. J., Kwon, K. K., de Donatis, G. M., Lee, J. H., Maurizi, M. R., & Kang, S. G. (2010). Crystal structure of Lon protease: Molecular architecture of gated entry to a sequestered degradation chamber. EMBO J, 29(20), 3520–3530

Chen, V. B., Arendall, W. B., Headd, J. J., Keedy, D. A., Immormino, R. M., Kapral, G. J., Murray, L. W., Richardson, J. S., & Richardson, D. C. (2010). MolProbity: All-atom structure validation for macromolecular crystallography. Acta Crystallogr D, 66(1), 12–21

Chen,. X, Zhang, S., Bi, F., Guo, C., Feng, L., Wang, H., Yao, H. & Lin, D. Crystal structure of the N domain of Lon protease from Mycobacterium avium complex. (2019) Protein Sci 28(9), 1720–1726.

de la Peña, A. H., Goodall, E. A., Gates, S. N., Lander, G. C., & Martin, A. (2018). Substrate-engaged 26S proteasome structures reveal mechanisms for ATP-hydrolysis-driven translocation. Science, 362(6418)

Deville, C., Carroni, M., Franke, K. B., Topf, M., Bukau, B., Mogk, A., & Saibil, H. R. (2017). Structural pathway of regulated substrate transfer and threading through an Hsp100 disaggregase. Sci Adv, 3(8), e1701726

Duman, R. E., & Löwe, J. (2010). Crystal structures of Bacillus subtilis Lon protease. J Mol Biol, 401(4), 653–670

Emsley, P., Lohkamp, B., Scott, W. G., & Cowtan, K. (2010). Features and development of Coot. Acta Crystallogr D, 66(4), 486–501

Gates, S. N., Yokom, A. L., Lin, J. B., Jackrel, M. E., Rizo, A. N., Kendsersky, N. M., Buell, C. E., Sweeny, E. A., Mack, K. L., Chuang, E., Torrente, M. P., Su, M., Shorter, J., & Southworth, D. R. (2017). Ratchet-like polypeptide translocation mechanism of the AAA+ disaggregase Hsp104. Science, 357(6348), 273–279

Goddard, T. D., Huang, C. C., Meng, E. C., Pettersen, E. F., Couch, G. S., Morris, J. H., & Ferrin, T. E. (2018). UCSF ChimeraX: Meeting modern challenges in visualization and analysis. Protein Sci, 27(1), 14–25

He, L., Luo, D., Yang, F., Li, C., Zhang, X., Deng, H., & Zhang, J.-R. (2018). Multiple domains of bacterial and human Lon proteases define substrate selectivity. Emerg Microbes Infec, 7(1), 149

Jakobi, A. J., Wilmanns, M., & Sachse, C. (2017). Model-based local density sharpening of cryo-EM maps. ELife, 6

Vaca Jacome, A.S., Rabilloud, T., Schaeffer-Reiss, C., Rompais, M., Ayoub, D., Lane, L., Bairoch, A., Van Dorsselaer, A, & Carapito, C. (2015). N-terminome analysis of the human mitochondrial proteome. Proteomics 15(14), 2519–2524.

Kereïche, S., Kováčik, L., Bednár, J., Pevala, V., Kunová, N., Ondrovičová, G., Bauer, J., Ambro, Ľ., Bellová, J., Kutejová, E., & Raška, I. (2016). The N-terminal domain plays a crucial role in the structure of a full-length human mitochondrial Lon protease. Sci Rep, 6, 33631

Lee, J., Feldman, A. R., Delmas, B., & Paetzel, M. (2007). Crystal structure of the VP4 protease from infectious pancreatic necrosis virus reveals the acyl-enzyme complex for an intermolecular self-cleavage reaction. J Biol Chem, 282(34), 24928–24937

Lee, S., Augustin, S., Tatsuta, T., Gerdes, F., Langer, T., & Tsai, F. T. F. (2011). Electron cryomicroscopy structure of a membrane-anchored mitochondrial AAA protease. J Biol Chem, 286(6), 4404–4411

Li, M., Gustchina, A., Rasulova, F. S., Melnikov, E. E., Maurizi, M. R., Rotanova, T. v, Dauter, Z., & Wlodawer, A. (2010). Structure of the N-terminal fragment of Escherichia coli Lon protease. Acta Crystallogr D, 66(Pt 8), 865–873

Li, M., Rasulova, F., Melnikov, E. E., Rotanova, T. v., Gustchina, A., Maurizi, M. R., & Wlodawer, A. (2005). Crystal structure of the N-terminal domain of E. coli Lon protease. Protein Sci, 14(11), 2895–2900

Liebschner, D., Afonine, P. v., Baker, M. L., Bunkoczi, G., Chen, V. B., Croll, T. I., Hintze, B., Hung, L. W., Jain, S., McCoy, A. J., Moriarty, N. W., Oeffner, R. D., Poon, B. K., Prisant, M. G., Read, R. J., Richardson, J. S., Richardson, D. C., Sammito, M. D., Sobolev, O. v., … Adams, P. D. (2019). Macromolecular structure determination using X-rays, neutrons and electrons: Recent developments in Phenix. Acta Crystallogr D, 75(Pt 10), 861–877

Lin, C. C., Su, S. C., Su, M. Y., Liang, P. H., Feng, C. C., Wu, S. H., & Chang, C. I. (2016). Structural Insights into the Allosteric Operation of the Lon AAA+ Protease. Structure, 24(5), 667–675

Lu, B., Lee, J., Nie, X., Li, M., Morozov, Y. I., Venkatesh, S., Bogenhagen, D. F., Temiakov, D., & Suzuki, C. K. (2013). Phosphorylation of human TFAM in mitochondria impairs DNA binding and promotes degradation by the AAA+ Lon protease. Mol Cell, 49(1), 121–132

Monroe, N., Han, H., Shen, P. S., Sundquist, W. I., & Hill, C. P. (2017). Structural basis of protein translocation by the Vps4-Vta1 AAA ATPase. ELife, 6

Münch, C., & Harper, J. W. (2016). Mitochondrial unfolded protein response controls matrix pre-RNA processing and translation. Nature, 534(7609), 710–713

Murshudov, G. N., Skubák, P., Lebedev, A. A., Pannu, N. S., Steiner, R. A., Nicholls, R. A., Winn, M. D., Long, F., & Vagin, A. A. (2011). REFMAC5 for the refinement of macromolecular crystal structures. Acta Crystallogr D, 67(4), 355–367

Nicholls, R. A., Fischer, M., Mcnicholas, S., & Murshudov, G. N. (2014). Conformation-independent structural comparison of macromolecules with ProSMART. Acta Crystallogr D, 70(9), 2487–2499

Peter, B., Waddington, C. L., Oláhová, M., Sommerville, E. W., Hopton, S., Pyle, A., Champion, M., Ohlson, M., Siibak, T., Chrzanowska-Lightowlers, Z. M. A., Taylor, R. W., Falkenberg, M., & Lightowlers, R. N. (2018). Defective mitochondrial protease LonP1 can cause classical mitochondrial disease. Hum Mol Genet, 27(10), 1743–1753

Pinti, M., Gibellini, L., Nasi, M., de Biasi, S., Bortolotti, C. A., Iannone, A., & Cossarizza, A. (2016). Emerging role of Lon protease as a master regulator of mitochondrial functions. Biochim Biophys Acta - Bioenergetics, 1857(8), 1300–1306

Puchades, C., Rampello, A. J., Shin, M., Giuliano, C. J., Wiseman, R. L., Glynn, S. E., & Lander, G. C. (2017). Structure of the mitochondrial inner membrane AAA+ protease YME1 gives insight into substrate processing. Science, 358(6363)

Puchades, C., Sandate, C. R., & Lander, G. C. (2020). The molecular principles governing the activity and functional diversity of AAA+ proteins. Nat Rev Mol Cell Bio, 21(1), 43–58

Punjani, A., Rubinstein, J. L., Fleet, D. J., & Brubaker, M. A. (2017). CryoSPARC: Algorithms for rapid unsupervised cryo-EM structure determination. Nat Methods, 14(3), 290–296

Quirós, P. M., Español, Y., Acín-Pérez, R., Rodríguez, F., Bárcena, C., Watanabe, K., Calvo, E., Loureiro, M., Fernández-García, M. S., Fueyo, A., Vázquez, J., Enríquez, J. A., & López-Otín, C. (2014). ATP-Dependent Lon Protease Controls Tumor Bioenergetics by Reprogramming Mitochondrial Activity. Cell Rep, 8(2), 542–556

Reinhard, L., Sridhara, S., & Hällberg, B. M. (2017). The MRPP1/MRPP2 complex is a tRNA-maturation platform in human mitochondria. Nucleic Acids Res, 45(21), 12469–12480

Ripstein, Z. A., Huang, R., Augustyniak, R., Kay, L. E., & Rubinstein, J. L. (2017). Structure of a AAA+ unfoldase in the process of unfolding substrate. eLife, 6

Ripstein, Z. A., Vahidi, S., Houry, W. A., Rubinstein, J. L., & Kay, L. E. (2020). A processive rotary mechanism couples substrate unfolding and proteolysis in the ClpXP degradation machinery. ELife, 9

Rosenthal, P. B., & Henderson, R. (2003). Optimal determination of particle orientation, absolute hand, and contrast loss in single-particle electron cryomicroscopy. J Mol Biol, 333(4), 721–745

Sauer, R. T., & Baker, T. A. (2011). AAA+ Proteases: ATP-fueled machines of protein destruction. Annu Rev Biochem, 80, 587–612

Scheres, S. H. W. (2012). RELION: Implementation of a Bayesian approach to cryo-EM structure determination. J Struct Biol, 180(3), 519–530

Shin, M., Puchades, C., Asmita, A., Puri, N., Adjei, E., Wiseman, R. L., Karzai, A. W., & Lander, G. C. (2020). Structural basis for distinct operational modes and protease activation in AAA+ protease Lon. Sci Adv, 6(21), eaba8404

Shin, M., Watson, E. R., Song, A.S., Mindrebo, J.T., Novick, S. J., Griffin, P., Wiseman, R. L., & Lander, G. C. (2021). Structures of the human LONP1 protease reveal regulatory steps involved in protease activation. Nat com, 12, 3239

Strauss, K. A., Jinks, R. N., Puffenberger, E. G., Venkatesh, S., Singh, K., Cheng, I., Mikita, N., Thilagavathi, J., Lee, J., Sarafianos, S., Benkert, A., Koehler, A., Zhu, A., Trovillion, V., McGlincy, M., Morlet, T., Deardorff, M., Innes, A. M., Prasad, C., … Suzuki, C. K. (2015). CODAS Syndrome Is Associated with Mutations of LONP1, Encoding Mitochondrial AAA+ Lon Protease. Am J Hum Genet, 96(1), 121–135

Tegunov, D., & Cramer, P. (2019). Real-time cryo-electron microscopy data preprocessing with Warp. Nat Methods, 16(11), 1146–1152

Tzeng, S.R., Tseng, Y.C, Lin, C.C., Hsu, C.Y., Huang, S.J., Kuo, Y.T., Chang, C.I. (2021). Molecular insights into substrate recognition and discrimination by the N-terminal domain of Lon AAA+ protease. eLife, 10:e64056.

Vieux, E. F., Wohlever, M. L., Chen, J. Z., Sauer, R. T., & Baker, T. A. (2013). Distinct quaternary structures of the AAA+ Lon protease control substrate degradation. P Natl Acad Sci USA, 110(22), E2002–2008

Wohlever, M. L., Baker, T. A., & Sauer, R. T. (2014). Roles of the N domain of the AAA+ Lon protease in substrate recognition, allosteric regulation and chaperone activity. Mol Microbiol, 91(1), 66–78

Zhang, K., Li, S., Hsieh, K-Y., Su, S-C., Pintilie, G. D., Chiu, W., Chang, C-I. (2020). Molecular basis for the ATPase-powered substrate translocation by the Lon AAA+ protease BioRxiv, 2020.04.29.068361

Zivanov, J., Nakane, T., & Scheres, S. H. W. (2019). A Bayesian approach to beam-induced motion correction in cryo-EM single-particle analysis. IUCrJ, 6(Pt 1), 5–17

