## supplementary materials for "A dual allosteric pathway drives human mitochondrial Lon"

**A dual allosteric pathway drives human mitochondrial Lon**
**Supplemental material**

Genís Valentín Gesé[1†], Saba Shahzad[1,3†], Carlos Pardo-Hernández[2†], Anna Wramstedt[2], Maria Falkenberg[2], B. Martin Hällberg[1,4*]

[1] Department of Cell and Molecular Biology, Karolinska Institutet, Sweden

[2] Department of Medical Biochemistry and Cell Biology, University of Gothenburg, Sweden

[3] Current address: Department of Medical Biochemistry and Biophysics, Umeå University, Sweden

[4] Centre for Structural Systems Biology (CSSB) and Karolinska Institutet VR-RÅC, Germany

[†]Joint First Authors

*Correspondence to:
B. Martin Hällberg.

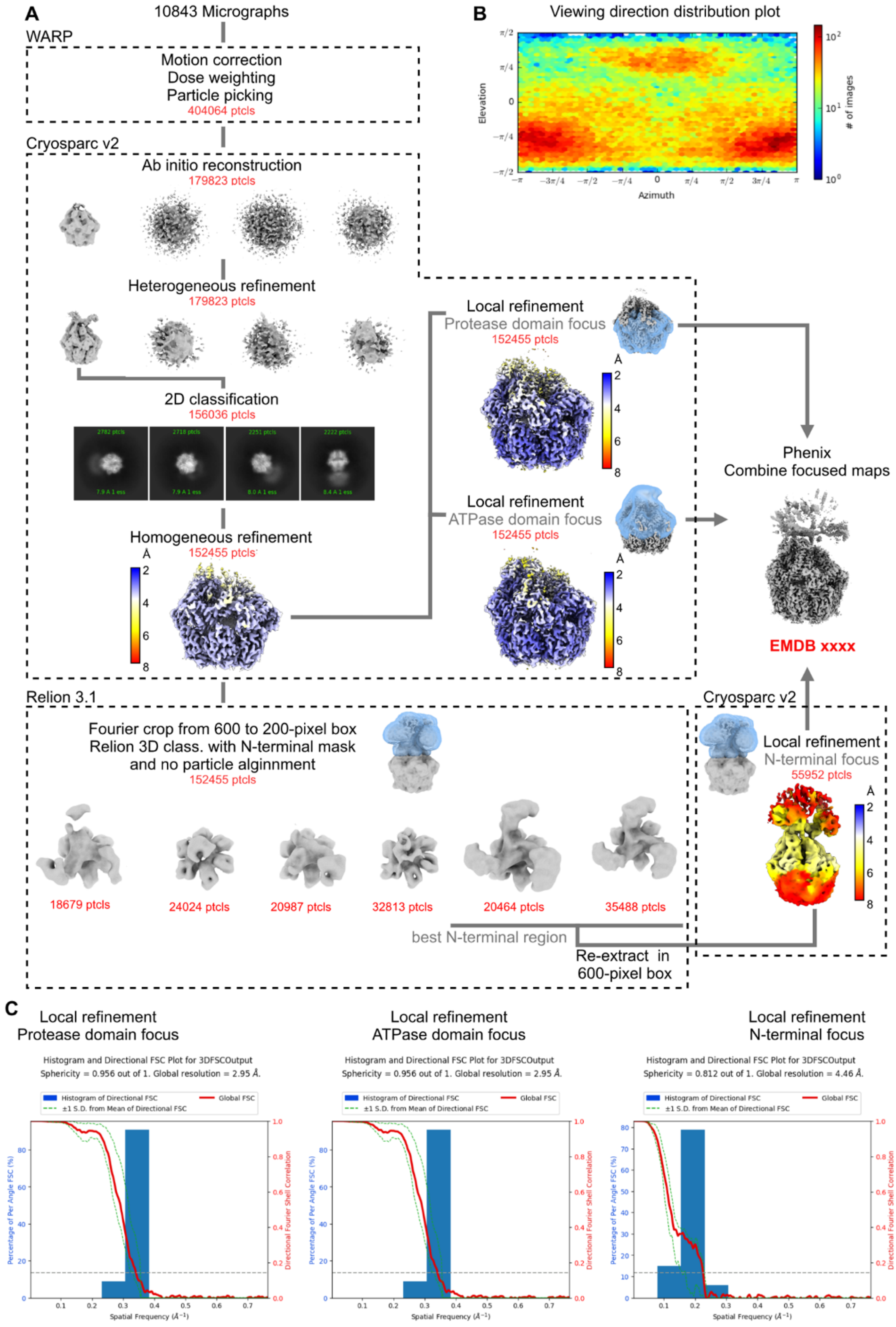


**Figure S1**. (**A**) Data processing strategy for Lon$^{SE}$. (**B)** Viewing distribution plot obtained from Cryosparc v2.15 for the ATPase-focused local refinement map. (**C**) 3DFSC (Zi Tan et al., 2017) histogram for the local refinement maps.

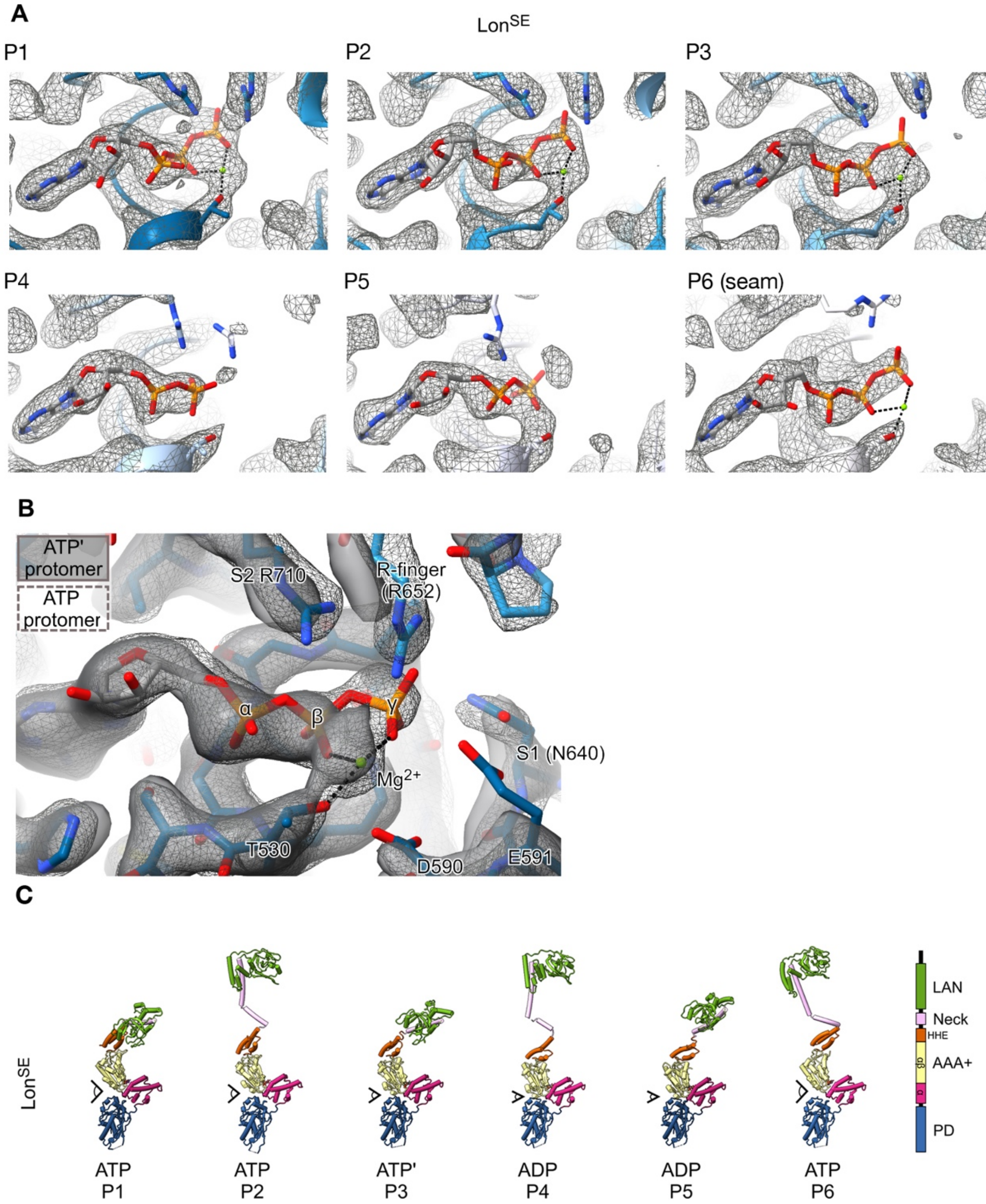


**Figure S2.** Nucleotide load analysis of Lon$^{SE}$. (**A**) Nucleotide density in Lon$^{SE}$. The protomers are indicated by numbers P1 to P6. (**B**) Comparison of the nucleotide density between P1 (ATP) and P3 (ATP'), shown as a mesh and a semi-transparent surface, respectively. The densities were superimposed by matching the P-loop of P1 and P3. The P1 ATP and the surrounding residues are shown in stick representation. (**C**) Individual representation of Lon$^{SE}$ protomers P1 to P6. The protomers (P1 to P6) are coloured by domain, as indicated in the legend on the right. The nucleotide is shown in stick representation. α, AAA+ α subdomain; αβ, AAA+ αβ subdomain; HHE, helical region. PS1βH, pre-sensor 1 β-hairpin.

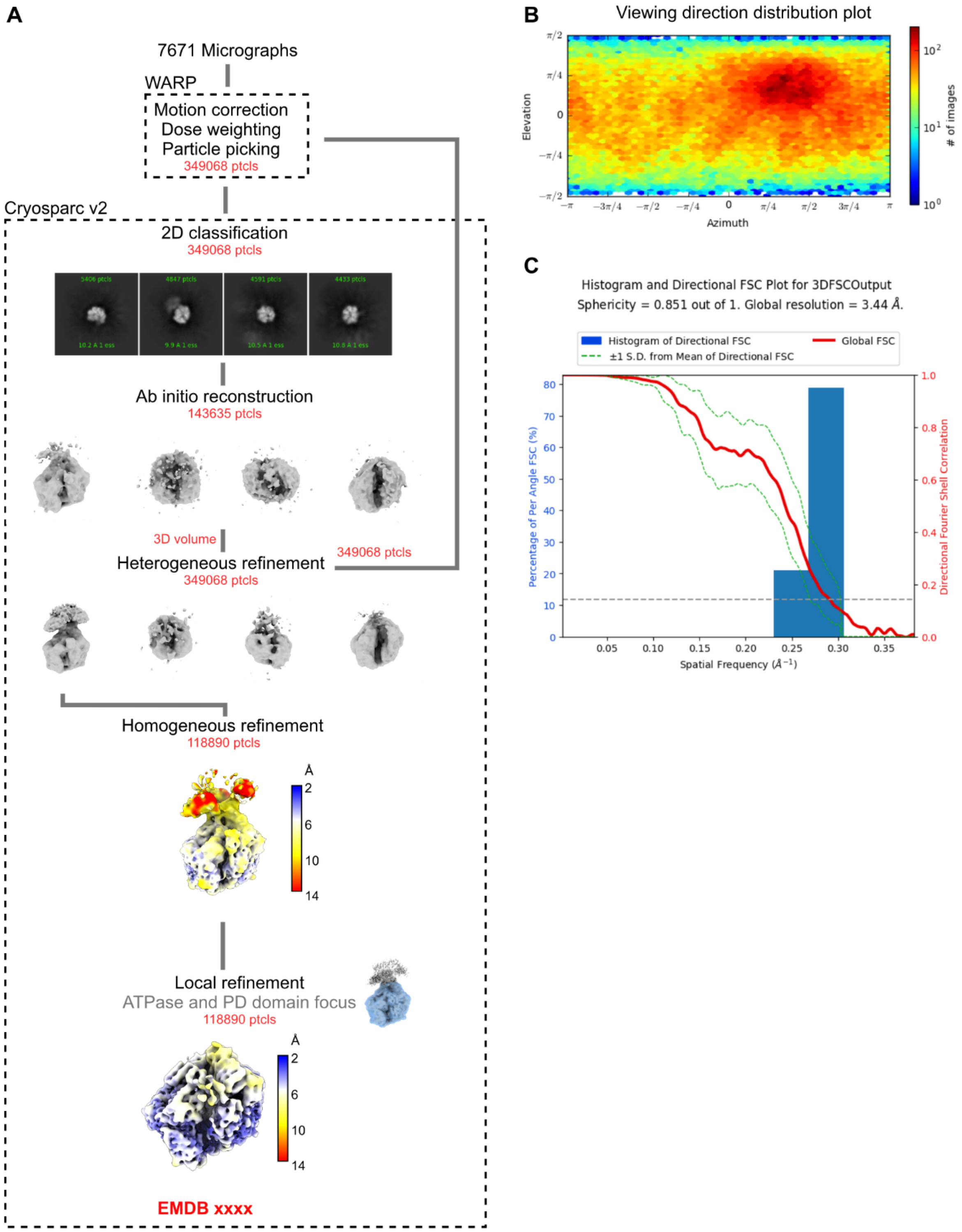


**Figure S3.** (A) Data processing strategy for Lon$^{Apo}$. (**B**) Viewing distribution plot obtained from Cryosparc v2.15 for the ATPase-focused local refinement map. (**C**) 3DFSC (Zi Tan et al., 2017) histogram for the local refinement map.

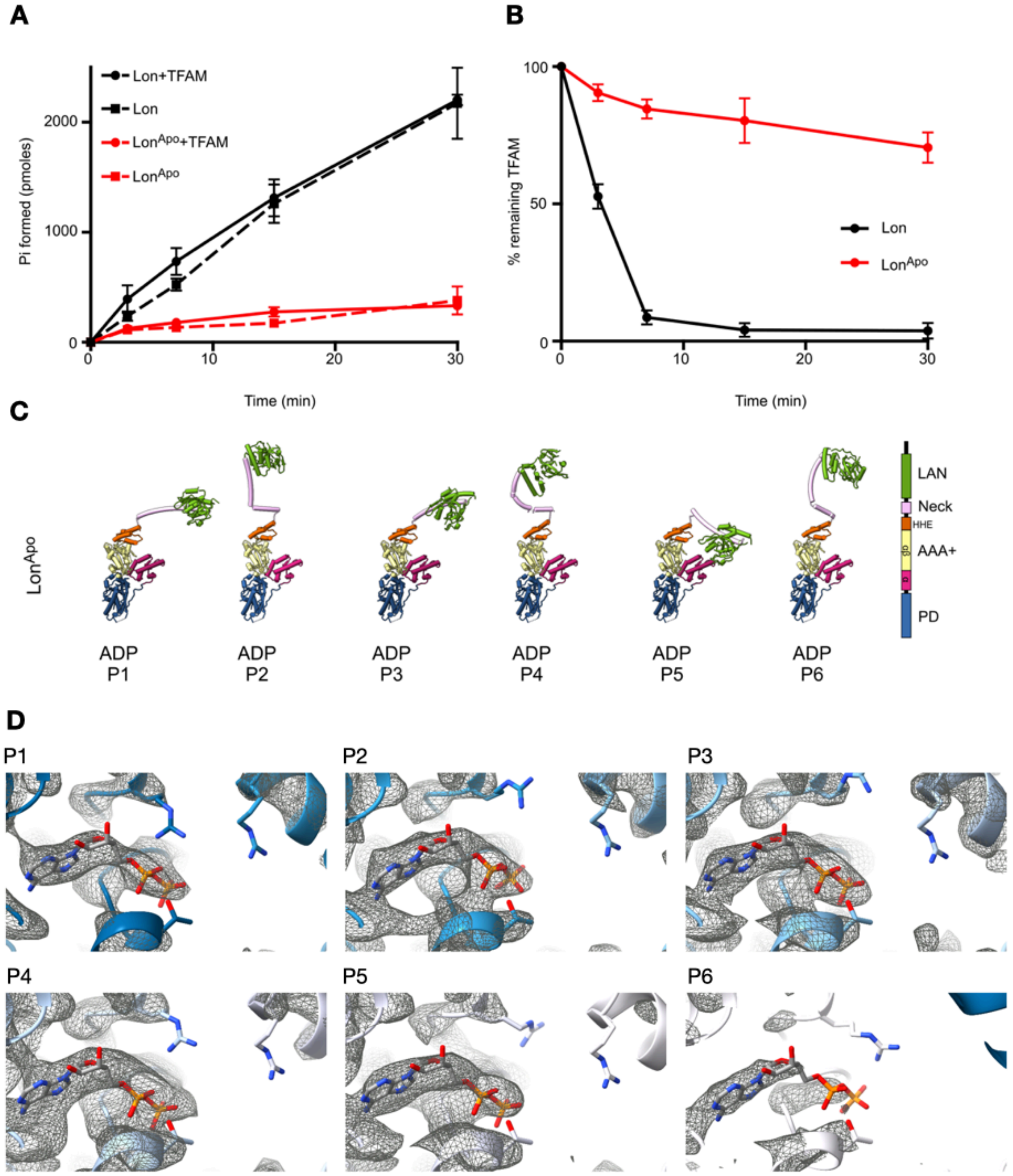


**Figure S4.** (**A**) Quantification of the ATP hydrolysis activity of Lon and Lon$^{Apo}$ over time (0-30 min). Solid line represents experiments done in the presence of TFAM, whereas dashed line represents experiments done in the absence of TFAM. Data is represented as mean ± standard deviation (n=3). (**B**) Quantification of the proteolytic activity of Lon and Lon$^{Apo}$ over time (0-30 min). Data is represented as mean ± standard deviation (n = 3). (**C**) Individual representation of Lon$^{Apo}$ P1 to P6. The protomers (P1 to P6) are coloured by domain, as indicated in the legend on the right. The nucleotide is shown in stick representation. $\alpha$, AAA+ $\alpha$ subdomain; $\alpha\beta$, AAA+ $\alpha\beta$ subdomain; HHE, helical region. PS1$\beta$H, pre-sensor 1 $\beta$-hairpin. (**D**) Nucleotide density in Lon$^{Apo}$. The protomers are indicated by P1 to P6.

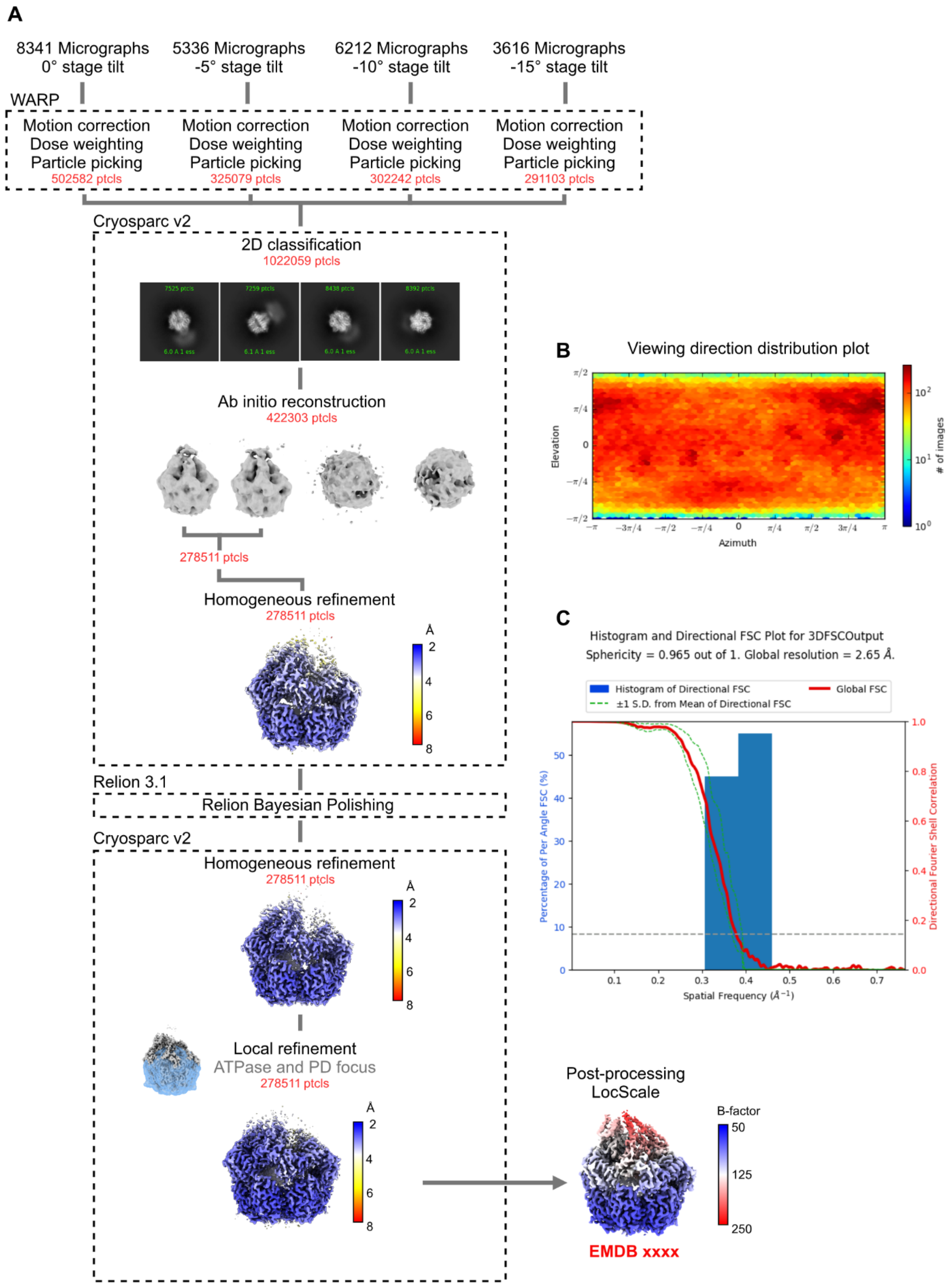


**Figure S5.** (**A**) Data processing strategy for Lon$^{K898A}$. (**B**) Viewing distribution plot obtained from Cryosparc v2.15 for the ATPase-focused local refinement map. (**C**) 3DFSC (Zi Tan et al., 2017) histogram for the local refinement map.

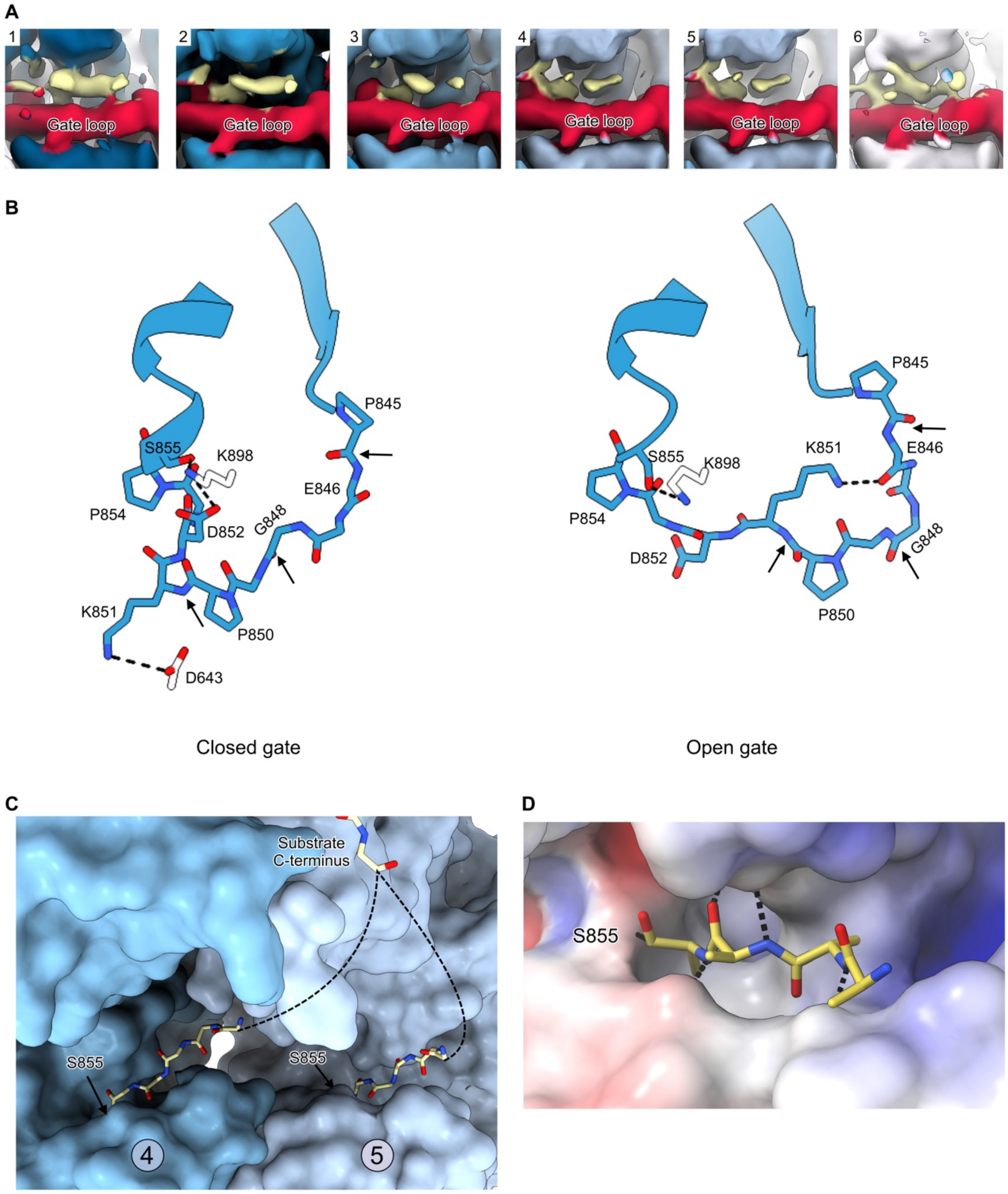


**Figure S6.** Substrate binding to the protease domains (**A**) The gate loop is in red and the substrate in yellow. All the panels are shown at the same map contour level. (**B**) Main chain atoms of the protease domain gate loop in the closed conformation ($Lon^{Apo}$, left panel) and the open conformation ($Lon^{SE}$, right panel). The side chain atoms for the residues stabilizing the loop conformations (P845, P850 and P854, as well as K851 and D852) are shown. The catalytic S855 and K898 (white) are shown. The black arrows indicate the peptide bonds which flip by 180°. (**C**) Possible connections between the substrate peptide (yellow sticks) observed in the AAA+ domain (top part) to the substrate peptides observed in the protease domains is shown by the dashed lines. The surface representation of Lon P4 and P5 is shown. (**D**) Surface electrostatic potential (-5 $-kTe^{-1}$, blue to +5 $-kTe^{-1}$, red) of the protease active site in $Lon^{K898A}$. The substrate is shown in yellow sticks. The dashed black lines indicate H-bonds.

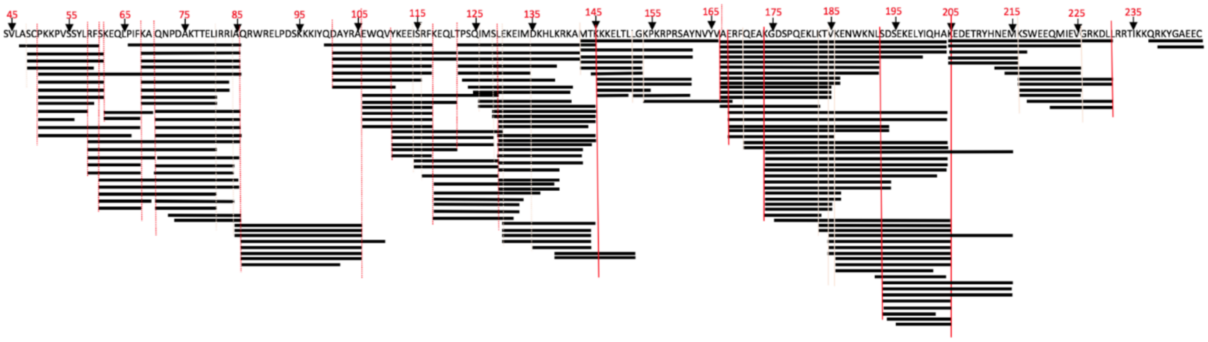


**Figure S7**. Peptides generated by the degradation of TFAM by Lon. Graphical representation of the TFAM peptides (horizontal black lines) produced by Lon hydrolysis as aligned to the mature TFAM sequence (44-246). The dotted red vertical lines highlight the cleaved peptide bond in the TFAM sequence.

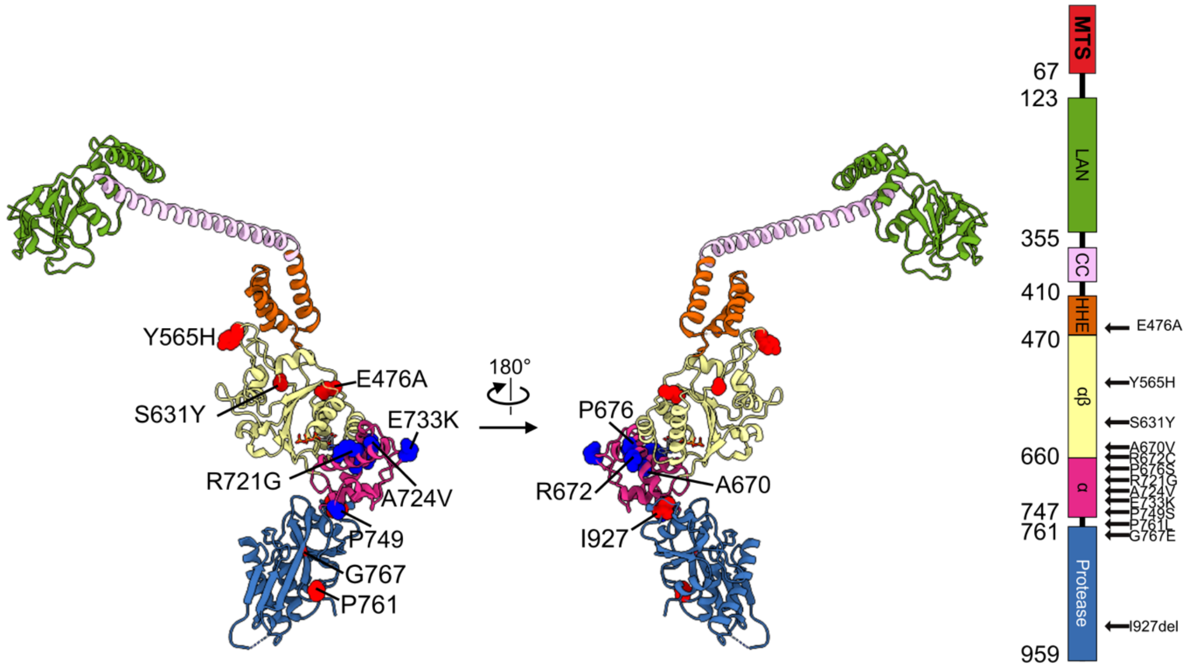


**Figure S8**. Clinically relevant mutations mapped on human Lon. The mutated residues are shown as sphere representation, either in red or blue for higher contrast with the background. α, AAA+ α subdomain; αβ, AAA+ αβ subdomain. HHE, helical region. MTS, mitochondrial targeting sequence. Mutations in the LonP1 gene were found to be the cause of Cerebral, Ocular, Dental, Auricular Skeletal (CODAS) syndrome, a complex multisystemic and developmental disorder. The natural variants that were found to be linked with CODAS (https://www.omim.org/entry/600373) mapped on the Lon structure show that four out of six reported mutations are on the α subdomain: P676S, R721G, A724V, E733K. The remaining two, S631Y and Y565H, are on the α/β-subdomain (Strauss et al., 2015). The P676S and R721G are on α21, whose N-terminus is directly involved in ATP binding while the C-terminus is involved in the surface interaction between the protomers. The A724V and E733K mutations are on the α15-helix that forms the major interface between the protomers so thereby potentially directly affecting the oligomerization. The S631Y mutation when first identified was believed to be present near the nucleotide binding pocket. But the present structure of Lon shows that it is present in the PS1βH (618-629). As reported here, main purpose of the PS1βH communicates the pore loop 1 with the ATP catalytic site via the ISS element. Therefore, we speculate that the mutation S631Y will abolish the substrate stimulation of ATP hydrolysis. The Y565H mutation is present in the pore loop 1 and directly contacts the substrate, explaining why this mutant is unable to bind and degrade substrates (Peter et al., 2018).

#### Table S1. Rate constants of TFAM degradation and end point ATP hydrolysis measurements

| | TFAM hydrolysis | | ATP hydrolysis (pmoles Pi formed after 30 minutes) | |
|---|---|---|---|---|
| | **Rate constant ($min^{-1}$)** | **%TFAM remaining after 30 minutes** | **+ TFAM** | **- TFAM** |
| **Lon res. 67-959** | 0.27 ± 0.05 | 3.8 ± 2.8 | 2202 ± 49 | 2174 ± 323 |
| **G106P, R563W and K594M** | 0.04 ± 0.01 | 70.5 ± 5.5 | 333 ± 6 | 381 ± 128 |
| **Y394A** | 0.053 ± 0.02 | 82.1 ± 0.3 | 340 ± 33 | 205 ± 33 |
| **L396A** | 0.066 ± 0.01 | 67 ± 1.1 | 386 ± 52 | 233 ± 28 |
| **Y599A** | 0.043 ± 0.01 | 42.9 ± 8.9 | 1151 ± 70 | 952 ± 16 |
| **E614A** | 0.039 ± 0.05 | 63.4 ± 5 | 1012 ± 183 | 573 ± 59 |

### Table S2. Cryo-EM data acquisition and image processing

| | | | | | |
|---|---|---|---|---|---|
| **Data collection** | | | | | |
| Sample | Lon 68-959 wild type + TFAM | | | Lon 68-959 K898A + TFAM | Lon 68-959 G106P, R563W, K594M + TFAM |
| Microscope | Krios G3i | | | Krios G3i | Krios G3i |
| Camera | Bioquantum K3 | | | Bioquantum K3 | Bioquantum K3 |
| Voltage (kV) | 300 | | | 300 | 300 |
| EFTEM SA Mag. | 130,000 | | | 130,000 | 130,000 |
| Calibrated physical pixel size (Å) | 0.654 | | | 0.654 | 0.654 |
| Fluency ($e^-$/Å$^2$) | 51 | | | 51 | 51 |
| Energy filter slit width (eV) | 10 | | | 10 | 10 |
| Number of frames | 60 | | | 60 | 60 |
| Underfocus range (□m) | 0.2-2.0 | | | 0.2-2.0 | 0.2-2.0 |
| **Image processing** | | | | | |
| Motion-correction software | Warp | | | Warp | Warp |
| Defocus-parameter estimation software | Warp | | | Warp | Warp |
| Particle-picking software | Warp | | | Warp | Warp |
| Micrographs used | 10,843 | | | 23,505 | 7,671 |
| Particle selected | 404,064 | | | 1,022,059 | 349,068 |
| 3D map classification and reconstruction software | CryoSPARC v2.15 and RELION 3.1 | | | CryoSPARC v2.15 and RELION 3.1 | CryoSPARC v2.15 |
| **EM maps** | **Lon$^{SE}$ N-terminal focus** | **Lon$^{SE}$ AAA+ focus** | **Lon$^{SE}$ PD focus** | **Lon$^{K898A}$** | **Lon$^{Apo}$** |
| EMDB code | | | | | |
| Particle images contributing to maps | 55,952 | 152,455 | 152,455 | 278,511 | 61,980 |
| Applied symmetry | C1 | C1 | C1 | C1 | C1 |
| Global resolution (FSC = 0.143, Å) | 7.42 | 2.70 | 2.75 | 2.65 | 4.11 |

EMDB, *Electron Microscopy Data Bank*; FSC, Fourier shell correlation

### Table S3. Model building and refinement statistics

| | | | |
|---|---|---|---|
| **Model building and refinement** | | | |
| Software | *Coot*, PHENIX Version 1.19-3660 | | |
| Sample | Lon$^{SE}$ | Lon$^{K898A}$ | Lon$^{Apo}$ |
| PDB code | XXXX | XXXX | XXXX |
| Number of residues (atoms) | 3,196 (25,091) | 3,220 (25,191) | 3,185 (25,002) |
| RMS bond length (outliers) | 0.002 (0) | 0.002 (0) | 0.002 (0) |
| RMS bond angle (outliers) | 0.438 (0) | 0.436 (1) | 0.601 (3) |
| Ramachandran outliers (%) | 0.00 | 0.00 | 0.00 |
| Ramachandran favoured (%) | 98.74 | 98.27 | 98.80 |
| Rotamer outliers (%) | 0.00 | 1.88 | 0.04 |
| C[illegible] outliers | 0.00 | 0.00 | 0.00 |
| Clashscore | 6.51 | 7.40 | 6.75 |
| MolProbity score | 1.36 | 1.62 | 1.37 |
| Refinement resolution (Å) | 2.7 | 2.5 | 2.9 |
| Map-Model CC_mask | 0.79 | 0.89 | 0.78 |
| Ligands | 2 ADP | 2 ADP | 6 ADP |
| | 4 ATP-$Mg^{2+}$ | 4 ATP-$Mg^{2+}$ | |
| B-factor (min/max/average in Å$^2$) | | | |
| protein | 90.91/231.06/125.67 | 63.33/230.20/103.71 | 206.94/661.58/330.08 |
| ligand | 93.67/156.88/116.95 | 87.70/136.77/103.16 | 255.34/384.72/309.80 |
| CaBLAM (%) | 0.54 | 0.48 | 0.86 |
| EM-ringer score | 3.35 | 2.62 | 0.52 |

CaBLAM, C-Alpha Based Low-resolution Annotation Method; CC, cross-correlation; PDB, Protein Data Bank; RMS, root-mean-square deviation

**Supplemental reference list**

Peter, B., Waddington, C. L., Oláhová, M., Sommerville, E. W., Hopton, S., Pyle, A., Champion, M., Ohlson, M., Siibak, T., Chrzanowska-Lightowlers, Z. M. A., Taylor, R. W., Falkenberg, M., & Lightowlers, R. N. (2018). Defective mitochondrial protease LonP1 can cause classical mitochondrial disease. *Human Molecular Genetics*, *27*(10), 1743–1753.

Strauss, K. A., Jinks, R. N., Puffenberger, E. G., Venkatesh, S., Singh, K., Cheng, I., Mikita, N., Thilagavathi, J., Lee, J., Sarafianos, S., Benkert, A., Koehler, A., Zhu, A., Trovillion, V., McGlincy, M., Morlet, T., Deardorff, M., Innes, A. M., Prasad, C., … Suzuki, C. K. (2015). CODAS Syndrome Is Associated with Mutations of LONP1, Encoding Mitochondrial AAA+ Lon Protease. *American Journal of Human Genetics*, *96*(1), 121–135.

Zi Tan, Y., Baldwin, P. R., Davis, J. H., Williamson, J. R., Potter, C. S., Carragher, B., & Lyumkis, D. (2017). Addressing preferred specimen orientation in single-particle cryo-EMthrough tilting. *Nature Methods*, *14*(8), 793–796.
